# Location-biased β-arrestin conformations direct GPCR signaling

**DOI:** 10.1101/2024.09.24.614742

**Authors:** Uyen Pham, Anand Chundi, Tomasz Maciej Stępniewski, Srikrishna Darbha, Dylan Scott Eiger, Sonia Gazula, Julia Gardner, Chloe Hicks, Jana Selent, Sudarshan Rajagopal

**Affiliations:** Department of Biochemistry, Duke University School of Medicine, Durham, NC 27710, USA; Department of Medicine, Duke University Medical Center, Durham, NC 27710, USA; Research Programme on Biomedical Informatics (GRIB), Department of Experimental and Health Sciences of Pompeu Fabra University (UPF)-Hospital del Mar Medical Research Institute (IMIM), 08003 Barcelona, Spain; InterAx Biotech AG, PARK InnovAARE, 5234 Villigen, Switzerland; Trinity College, Duke University, Durham, NC, 27710, USA; Department of Medicine, Brigham and Women’s Hospital, Boston, MA, 02215, USA; Harvard Medical School, Boston, MA, 02215, USA; Penn Dental Medicine, University of Pennsylvania, Philadelphia, PA, 19104, USA; Perelman School of Medicine, University of Pennsylvania, Philadelphia, PA, 19104, USA; Yale University School of Medicine, New Haven, CT, 06510, USA

**Keywords:** beta-arrestin, GPCR, G protein-coupled receptor, location bias, biased agonism, biased signaling, biosensors, conformations, FlAsH, catalytic activation, MAP kinase

## Abstract

β-arrestins are multifunctional intracellular proteins that regulate the desensitization, internalization and signaling of over 800 different G protein-coupled receptors (GPCRs) and interact with a diverse array of cellular partners^1,2^. Beyond the plasma membrane, GPCRs can initiate unique signaling cascades from various subcellular locations, a phenomenon known as “location bias”^3,4^. Here, we investigate how β-arrestins direct location-biased signaling of the angiotensin II type I receptor (AT1R). Using novel bioluminescence resonance energy transfer (BRET) conformational biosensors and extracellular signal-regulated kinase (ERK) activity reporters, we reveal that in response to the endogenous agonist Angiotensin II and the β-arrestin-biased agonist TRV023, β-arrestin 1 and β-arrestin 2 adopt distinct conformations across different subcellular locations, which are intricately linked to differential ERK activation profiles. We also uncover a population of receptor-free catalytically activated β-arrestins in the plasma membrane that exhibits insensitivity to different agonists and promotes ERK activation on the plasma membrane independent of G proteins. These findings deepen our understanding of GPCR signaling complexity and also highlight the nuanced roles of β-arrestins beyond traditional G protein pathways.

## INTRODUCTION

G protein-coupled receptors (GPCRs) are one of the largest and most versatile families of cell surface receptors, playing a crucial role in converting extracellular stimuli into intracellular signals^5^. Beyond the classical model of GPCR signaling, the concept of “biased signaling” has opened up new dimensions in our understanding of GPCR function^6^. Biased signaling occurs when different ligands, despite binding to the same GPCR, stabilize the receptor in distinct conformations that preferentially activate specific downstream pathways—either through G proteins or β-arrestins. This phenomenon is not only ligand-dependent but also influenced by other factors such as the specific transducer isoform involved, transducer conformation, cell type, and subcellular localization^7^. Recent studies have highlighted the importance of this spatial aspect, known as “location bias”, in which the differential signaling outputs that arise depend on where in the cell the receptor is located. While GPCRs were once thought to signal exclusively from the PM, it is now evident that they can continue to signal from various intracellular compartments^3^, including endosomes^8–11^, the Golgi apparatus^12,13^, nucleus^14,15^, and mitochondria^16,17^. This subcellular compartmentalization of signaling pathways adds another layer of regulatory complexity, contributing to the specificity and diversity of cellular responses.

Despite these advances, our understanding of the mechanisms underlying location bias and how β-arrestins contribute to this remains incomplete. Although initially discovered as desensitizers of G protein-mediated signaling, β-arrestins have emerged as critical modulators of GPCR signaling, capable of directing receptor trafficking and signaling depending on their interactions with the receptor^1,2,18^. Recent studies demonstrated that in addition to the canonical trafficking of β-arrestins associated with the receptor, catalytic activation of β-arrestins by the receptor core can localize β-arrestins to the plasma membrane (PM) and endosomes without co-trafficking of the receptor^19,20^. Structural studies have provided snapshots of how biased ligands can stabilize GPCRs and β-arrestins in active conformations, leading to selective recruitment of downstream effectors^21–26^. Additionally, different structures and phosphorylation patterns of the receptor induced by different ligands have been shown to promote distinct recruitment and active conformations of β-arrestins^27–32^. This dynamic structure-function relationship allows β-arrestins to selectively regulate different signaling pathways and modulate cellular functions with high specificity downstream of GPCRs. However, these studies often rely on purified systems that lack the dynamic context of living cells, where β-arrestins can adopt a range of conformations based on their environment.

In this study, we investigated the relationship between β-arrestin and signaling in different subcellular locations using novel conformational biosensors for β-arrestin 1 and β-arrestin 2, using Fluorescent Arsenical Hairpin (FlAsH) probes that complement in different subcellular locations (NanoBiT FlAsH). These NanoBiT FlAsH biosensors allowed us to monitor β-arrestin conformations in real-time across various subcellular compartments in living cells. At the angiotensin II type 1 receptor (AT1R), we found that β-arrestins adopt distinct, location-specific, and agonist-specific conformations at the receptor, the PM or in early endosomes. Additionally, we uncovered a population of catalytically activated β-arrestins at the PM which exhibit unique conformational states that are insensitive to ligand bias and is capable of promoting extracellular signal-regulated kinase (ERK) activation independently of G proteins. Furthermore, we demonstrated biased activation of subcellular ERK signaling cascades that were differentially regulated by G proteins, β-arrestins, and receptor endocytosis. By elucidating the relationship between ligand bias, β-arrestin conformation, and subcellular localization, we provide new insights into the spatial dynamics of GPCR signaling.

## RESULTS

### AT1R biased agonists promote distinct recruitment patterns of β-arrestins 1 and 2 at the receptor, plasma membrane, and endosomes

We first determined whether endogenous (AngII) and β-arrestin-biased (TRV023) AT1R agonists would induce different subcellular localization patterns of β-arrestins 1 and 2. Using a split luciferase complementation system (NanoBiT)^33^, we monitored the trafficking of SmBiT-β-arrestin 1 or SmBiT-β-arrestin 2 to the receptor (AT1R-LgBiT), a PM marker (LgBiT-CAAX), or an early endosome marker (2x-FYVE-LgBiT) in HEK293T cells expressing FLAG-AT1R (**Figure 1a-c**). Cells were stimulated with either the endogenous agonist angiotensin II (AngII) or the β-arrestin-biased agonist TRV120023 (TRV023). AngII induced robust recruitment of both β-arrestin isoforms to the AT1R with higher levels of β-arrestin 2 recruitment (**Figure 1d**). Consistent with previous studies, both β-arrestin isoforms were recruited to the AT1R with similar ligand potencies^34^. A similar pattern was observed with TRV023 although with reduced potency and efficacy. At the PM, β-arrestin 2 was trafficked significantly more than β-arrestin 1, although, unlike the recruitment pattern at the receptor, AngII and TRV023 displayed no measurable difference in their EC_50_ (**Figure 1e**). Interestingly, β-arrestin 2 was internalized into early endosomes with a significant difference in potency and efficacy between AngII and TRV023, while there was markedly less internalization of β-arrestin 1 (**Figure 1f**). Using confocal microscopy, we also observed consistent patterns of β-arrestin trafficking to the PM and endosomes following stimulation with AngII and TRV023 (**Extended Data Figure 1**). The Emax and EC_50_ values are summarized in **Table 1**.

**Figure 1:**
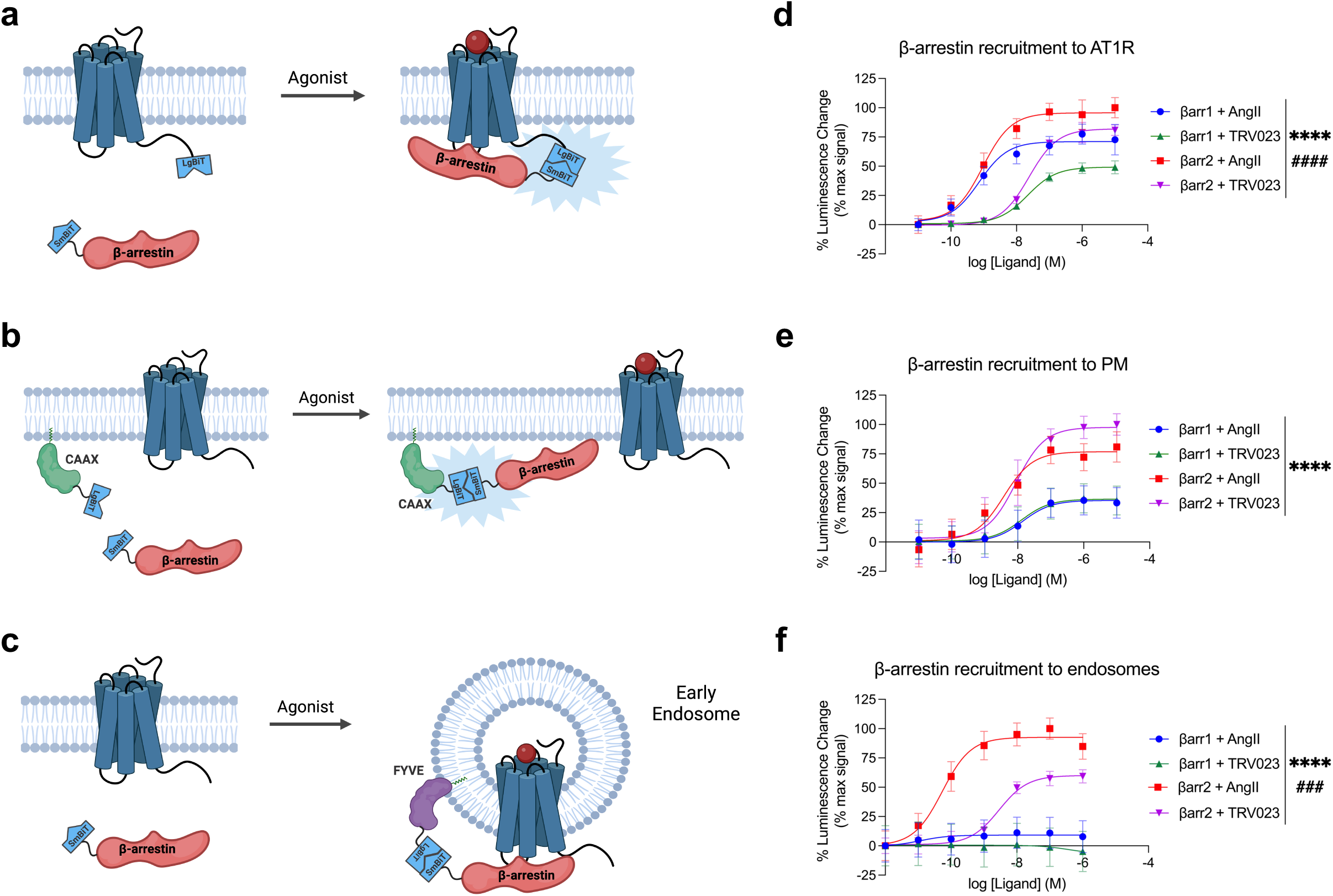
AngII and TRV023 promote different trafficking patterns of β-arrestins 1 and 2. (**a-c**) Schematics of NanoBiT assay monitoring the trafficking of β-arrestin 1 or β-arrestin 2 to the AT1R, PM, and early endosomes. **(d-f)** Dose response curves of the trafficking of β-arrestin isoforms to the AT1R (d), PM (e), and early endosomes (f). HEK293T cells were stimulated with agonist at the concentrations listed. Data is shown as percent change over vehicle normalized to max signal. Data represents mean ± SEM of *n* independent biological replicates, n=3 for AT1R and endosomes, n=5 for PM. One-way ANOVA with Tukey’s multiple comparison test to compare the Emax and EC50 values. * denotes the statistically significant differences between the Emax values. ^#^ denotes the statistically significant differences between the EC50 values. ****P<0.0001; ^###^P<0.0005; ^####^P<0.0001.

**Table 1.** – Emax and EC_50_ of β-arrestin recruitment.

| Location | β-arrestin Isoform/Ligand | E<sub>max</sub><br>(% max signal) | logEC<sub>50</sub> (M) |
| --- | --- | --- | --- |
| **AT1R** | β-arrestin1 – AngII | 71.06 ± 2.88<sup>***/###</sup> | -9.13 ± 0.16<sup>ns/####</sup> |
| | β-arrestin1 – TRV023 | 49.26 ± 0.56<sup>****/###</sup> | -7.66 ± 0.03<sup>ns/####</sup> |
| | β-arrestin2 – AngII | 95.65 ± 2.61<sup>***/##</sup> | -9.03 ± 0.11<sup>ns/####</sup> |
| | β-arrestin2 – TRV023 | 81.97 ± 1.67<sup>****/##</sup> | -7.62 ± 0.06<sup>ns/####</sup> |
| **Plasma**<br>**Membrane** | β-arrestin1 – AngII | 35.39 ± 1.45<sup>****/ns</sup> | -7.85 ± 0.12<sup>ns/ns</sup> |
| | β-arrestin1 – TRV023 | 36.49 ± 1.13<sup>****/ns</sup> | -7.89 ± 0.10<sup>ns/ns</sup> |
| | β-arrestin2 – AngII | 76.87 ± 4.71<sup>****/##</sup> | -8.42 ± 0.22<sup>ns/ns</sup> |
| | β-arrestin2 – TRV023 | 97.63 ± 4.32<sup>****/##</sup> | -8.05 ± 0.14<sup>ns/ns</sup> |
| **Early**<br>**Endosome** | β-arrestin1 – AngII | 9.16 ± 0.98<sup>****/#</sup> | -10.94 ± 0.50<sup>ns/###</sup> |
| | β-arrestin1 – TRV023 | -6.81 ± 3.95<sup>****/#</sup> | -6.44 ± 0.72<sup>*/###</sup> |
| | β-arrestin2 – AngII | 93.92 ± 3.60<sup>****/###</sup> | -10.14 ± 0.15<sup>ns/ns</sup> |
| | β-arrestin2 – TRV023 | 60.05 ± 1.82<sup>****/###</sup> | -8.54 ± 0.09<sup>*/ns</sup> |

The values were obtained from the nonlinear fit of the dose-response curves of β-arrestin recruitment.
Data represents mean ± SEM of *n* independent biological replicates, *n*=3 for AT1R and endosomes, *n*=5 for PM. One-way ANOVA with Tukey's multiple comparison test to compare the E<sub>max</sub> and EC<sub>50</sub> values.
\* denotes the statistically significant differences between the β-arrestin isoforms for a specific ligand. # denotes the statistically significant differences between the AT1R ligands for a specific isoform. \*P<0.05; \*\*\*P<0.0005; \*\*\*\*P<0.0001; #P<0.05; ##P<0.005; ###P<0.0005; ####P<0.0001; ns, not significant.

**Table 2.**
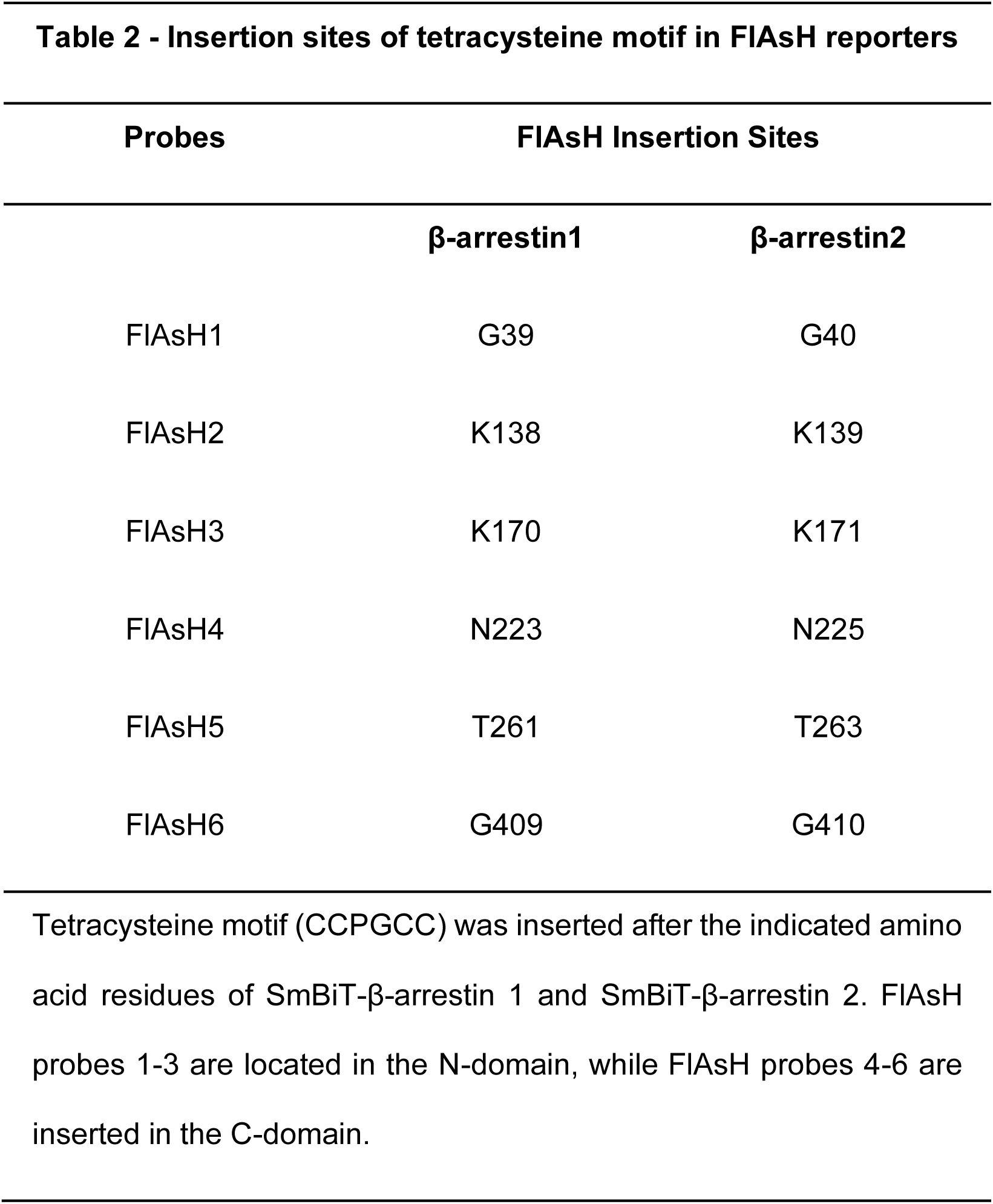
– Insertion sites of tetracysteine motif in FlAsH reporters.

| Probes | FIAsh Insertion Sites |  |
| --- | --- | --- |
| | β-arrestin1 | β-arrestin2 |
| FIAsh1 | G39 | G40 |
| FIAsh2 | K138 | K139 |
| FIAsh3 | K170 | K171 |
| FIAsh4 | N223 | N225 |
| FIAsh5 | T261 | T263 |
| FIAsh6 | G409 | G410 |

Tetracysteine motif (CCPGCC) was inserted after the indicated amino acid residues of SmBiT-β-arrestin 1 and SmBiT-β-arrestin 2. FIAsh probes 1-3 are located in the N-domain, while FIAsh probes 4-6 are inserted in the C-domain.

### AT1R biased agonists promote distinct conformations of β-arrestins 1 and 2 at the receptor and the endosomes but not the plasma membrane

As different conformations of β-arrestins are associated with distinct signaling profiles, such as GPCR desensitization, endocytosis, and MAPK activation^9,19,35–37^, we hypothesized that β-arrestins adopt different conformations in different locations. To determine these location-dependent conformations, we developed a Bioluminescence Resonance Energy Transfer (BRET)-based conformational biosensor called NanoBiT FlAsH, which couples components of the NanoBiT complementation system with a previously described intramolecular FlAsH BRET assay^9,27,33^. We generated six SmBiT-β-arrestin1-FlAsH probes and six SmBiT-β-arrestin2-FlAsH probes containing a tetracysteine motif (CCPGCC), which forms a hairpin loop that can bind to the fluorescent organoarsenic dye FlAsH-EDT2 with high affinity (**Figure 2a**). When a FlAsH sensor is recruited to the LgBiT-tagged location marker, their complementation forms a fully functional NanoLuciferase (NLuc), which emits a bioluminescence signal. The luminescence signal undergoes resonance energy transfer with the FlAsH acceptor located at different sites within β-arrestin, thereby generating a BRET signal that reports on β-arrestin conformation (**Figure 2b**). All of the SmBiT-β-arrestin-FlAsH probes were recruited to the receptor, the PM, and the endosomes after agonist stimulation, demonstrating that the tetracysteine motifs do not prevent β-arrestin recruitment (**Extended Data Figure 2**). Changes in the intramolecular BRET signals from six distinct regions within β-arrestin (FlAsH1-6) following AT1R stimulation were visualized using radar plots. In each plot, the shape of each six-sided figure represents the overall conformational signatures of β-arrestin at different subcellular locations.

**Figure 2:**
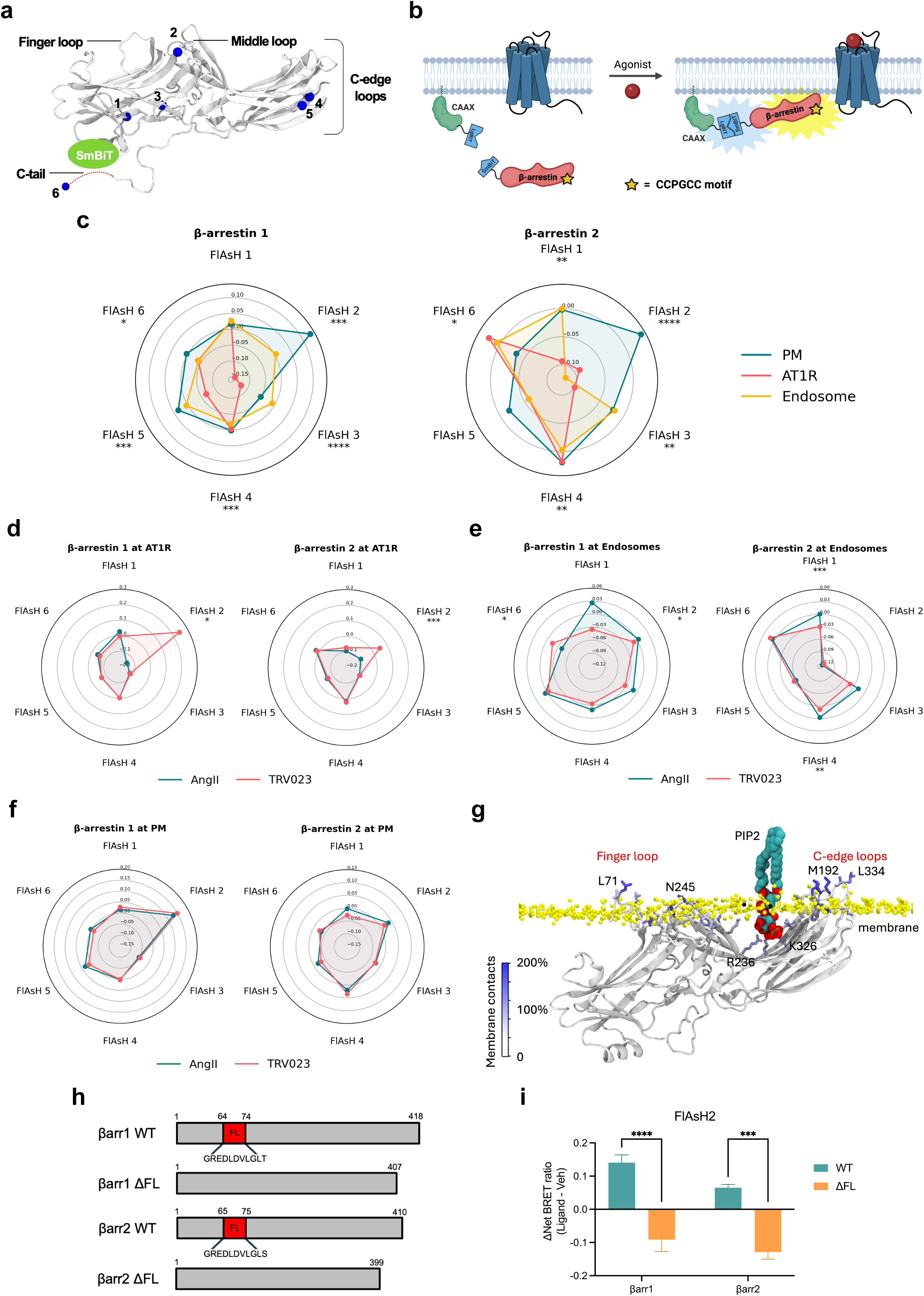
AT1R agonists promote distinct conformations of β-arrestins 1 and 2 at the receptor and endosomes but not the plasma membrane. **(a)** Schematic of NanoBiT FLAsH conformational biosensors. The tetracysteine motif CCPGCC (blue) is inserted after amino acid G39, K138, K170, N223, T261 and G409 for SmBiT-β-arrestin 1 or G40, K139, K171, N225, T263 and G410 for SmBiT-β-arrestin 2 to generate FlAsH 1-6, respectively. **(b)** Diagram of the NanoBiT FlAsH assay to detect the conformations of β-arrestins 1 and 2 at the receptor, the PM, or early endosomes. **(c)** Radar plots of the BRET signals from six FlAsH probes represent the location-specific conformations of β-arrestin 1 and β-arrestin 2 following stimulation with 1 μM AngII at the AT1R, PM or endosomes. Data represents mean ± SEM of *n* independent biological replicates. For AT1R, FlAsH 1: n=3, FlAsH 2,4: n=4, FlAsH 3, 5, 6: n=5. For CAAX, FlAsH 1-4: n=4, FlAsH 5: n=5, FlAsH 6: n=6. For 2xFYVE, FlAsH 1-5: n=4, FlAsH 6: n=3. One-way ANOVA with Tukey’s post hoc test comparing different subcellular locations for a specific FlAsH probe. *P<0.05; **P<0.005; ***P<0.0005; ****P<0.0001. **(d, e, f)** Radar plots comparing the ligand-specific effects on the conformational profiles of β-arrestins 1 and 2 in different subcellular locations. Cells were stimulated with 1 μM AngII or 10 μM TRV023. Data represents mean ± SEM of *n* independent biological replicates. Data with TRV023 has the same number of replicates as AngII, except FlAsH 5 (CAAX): n=7 and FlAsH 2 (FYVE): n=5. Unpaired two-tailed t-tests comparing AngII vs. TRV023 for each FlAsH sensor. *P<0.05; **P<0.005; ***P<0.0005; ****P<0.0001. **(g)** MD simulations of membrane-bound β-arrestin 1 anchored to the lipid bilayer with the finger loop, C-loop, and C-edge loops (3 x 500 ns). Residues have been colored according to the stability of contacts formed with the membrane (the frequency of contacts of each residue with individual membrane components has been calculated and summed in a per-residue fashion; residues with higher values form more stable interactions with the membrane). The position of PIP2 is also highlighted. **(h)** Diagram of WT β-arrestin 1, WT β-arrestin 2, and their finger loop deletion mutants. The position of the finger loop is highlighted in red. **(i)** Changes in FlAsH 2 signal of PM-localized β-arrestin 1 and β-arrestin 2 with deletion of the finger loop region. Data represents mean ± SEM, n=4 independent biological replicates. Two-way ANOVA with Šídák’s multiple comparisons comparing WT vs. ΔFL mutants. ***P<0.0005; ****P<0.0001.

First, we found that both β-arrestin isoforms displayed distinct location-specific conformations at the AT1R, PM, and early endosomes upon AngII stimulation (**Figure 2c**), with different kinetic profiles for each NanoBiT FlAsH reporter (**Extended Data Figure 3**). Significant differences in the conformational states were found in important regions in β-arrestins, particularly of the middle loop (FlAsH 2), C-edge loops (FlAsH 4 and FlAsH 5), and C-terminal tail (FlAsH 6). The β-arrestin-biased ligand TRV023 induced β-arrestin conformational signatures distinct from those promoted by AngII when β-arrestin was localized to the receptor and endosomes (**Figure 2d, e**). FlAsH 4 has previously been shown to report on the interdomain rotation between the N– and C-domain associated with β-arrestin activation^9,38^. Our data showed that the FlAsH 4 signal at the receptor did not differ between β-arrestins 1 and 2 or between AngII and TRV023 stimulation, suggesting that the β-arrestin isoforms underwent a similar degree of interdomain twist in response to different ligands.

β-arrestins bind GPCRs in two main modes: in the ‘tail’ conformations, they primarily interact with the phosphorylated C-terminus of the receptor, while in the ‘core’ conformation, they engage the receptor’s core using the finger loop in addition to the C-tail interaction^24,39^. The middle loop (FlAsH 2), which directly interacts with the central crest of the receptor in the core conformation, showed significant ligand-specific differences for both β-arrestins 1 and 2 at the receptor (**Figure 2d**). This is consistent with a model based on the structure of neurotensin receptor 1 in complex with β-arrestin 1 (PDB code: 6UP7), which suggests that the FlAsH 2-bearing middle loop is likely buried within the central crest of the receptor, thus reducing its mobility and resulting in smaller signal observed with AngII stimulation (**Extended Data Figure 4a**)^40^. Meanwhile, when comparing the structure of inactive β-arrestin 1 with the structure of β-arrestin 1 bound to the phosphorylated peptide of human V2 vasopressin receptor (V2Rpp), the position relating to FlAsH 2 insertion was observed to move closer toward the NLuc donor at the N-terminus, which would lead to an increase in BRET signal consistent with TRV023 treatment (**Extended Data Figure 4b**)^41,42^. These models support that AngII primarily promotes a core conformation of β-arrestin at the receptor while TRV023 stabilizes a tail conformation.

In contrast to the ligand-dependent differences occurring at the receptor and endosomes, the AT1R agonists promoted nearly identical conformations of β-arrestins at the PM, consistent with a catalytically activated population insensitive to ligand bias previously observed in β-arrestin trafficking to the PM^19,43^ (**Figure 2f**). Previous experimental data suggests that β-arrestin can anchor within the membrane using both the C-edge and finger loop regions^43^. To establish whether the FlAsH signatures obtained for β-arrestin 1 and 2 relate to a fully anchored (with both finger loop and C-edge regions embedded in the membrane) or partially anchored β-arrestin, we carried out molecular dynamics (MD) simulations (3 x 500 ns) of membrane-bound β-arrestin 1. Our simulations supported the ability of β-arrestin 1 to anchor within the membrane using the finger loop, C-loop, and C-edge loops (**Figure 2g**). Several residues were identified to form direct contact with the membrane, including Leu71 in the finger loop, Asn245 in the C-loop, and Met192 and Leu334 in the C-edge loops. Interestingly, when comparing the simulations of membrane-bound β-arrestin 1 to the same protein in solution, we observed a marked reduction in the flexibility and mobility of the middle loop, resulting from the insertion of the finger loop into the PM (**Extended Data Figure 5a**). This compression of the middle loop toward the N-terminus is also supported by our experimental data, which showed a positive change in net BRET ratio for FlAsH 2 in both β-arrestin isoforms, strongly suggesting that the BRET readouts for β-arrestin 1 and 2 correspond to a fully anchored conformation.

The role of C-edge loops in anchoring β-arrestin to the PM has been previously demonstrated in several biochemical and structural studies^23,24,44^. To validate whether the finger loop can bind to the membrane, we utilized a mutagenesis approach to remove the finger loop region of β-arrestin (G64-T74 of β-arrestin 1, G65-S75 of β-arrestin 2) and assessed the conformational state of FlAsH 2 at the PM (**Figure 2h**). The ΔFL mutants were still able to recruit to the PM, although to a lesser extent than wild-type (WT) FlAsH 2 (**Extended Data Figure 5b**). Our data showed that the finger loop deletion significantly altered the conformation reported by FlAsH 2 for both β-arrestin isoforms from positive BRET to negative BRET values (**Figure 2i**). These BRET values represent a conformational change of the middle loop from a compressed state due to finger loop insertion into the PM, to a relaxed and flexible state when the N-domain of ΔFL mutants is released from the membrane, suggesting the importance of the finger loop in anchoring β-arrestin to the lipid bilayer.

These data provide strong evidence that the conformational profiles of β-arrestin 1 and 2 in living cells is heavily influenced by its subcellular location. Notably, no agonist-induced changes were observed in β-arrestin 1 and 2 conformations at the PM. β-arrestins have been previously shown to undergo catalytic activation by other receptors, allowing them to dissociate from the receptor and interact with the PM^19,20^. This data is consistent with the recruitment data (**Figure 1**), suggesting a distinct effect of different AT1R ligands on the initial recruitment of β-arrestin to the receptor and not the subsequent localization of catalytically activated β-arrestins to the PM. The difference in the EC_50_ of the AT1R ligands likely reflects the distinct interaction of β-arrestins with different ligand-bound receptor conformations at the membrane or in early endosomes, while the catalytically activated pool of β-arrestins at the PM is not in contact with the receptor. Overall, these data suggest that different AT1R ligands promote distinct trafficking patterns of receptor-bound β-arrestins at the membrane and endosomes, while the conformation of catalytically activated β-arrestins at the PM remains unaffected by the agonist-bound receptor.

### The conformations of catalytically active β-arrestins at the plasma membrane depend on different lipid microdomains

Given our previous observation that the pool of β-arrestins localized at the PM had differential trafficking patterns and conformational signatures compared to β-arrestins at the receptor, we hypothesized that catalytically activated β-arrestin pools at the PM adopt unique conformations in different lipid microdomains. Lipid rafts are small, dynamic, and self-assembled PM compartments enriched with sphingolipids, cholesterol and saturated phospholipids^45–47^. Several GPCRs, β-arrestins, and other effectors such as adenylyl cyclase are concentrated in these microdomains^47–49^. To test our hypothesis, we investigated the conformational profiles of β-arrestins 1 and 2 at non-raft membrane and lipid rafts using the NanoBiT FlAsH assay. We co-transfected FLAG-AT1R and the FlAsH probes with a prenylation sequence (LgBiT-CAAX) as a marker for non-raft membrane or a previously described consensus sequence for myristoylation and palmitoylation (MyrPalm-LgBiT) for lipid rafts^50–52^. Cells were then stimulated with 1 μM AngII or 10 μM TRV023. Consistent with our FlAsH data at the non-raft membrane, β-arrestins 1 and 2 adopted distinct isoform-specific conformations in lipid rafts, specifically at the middle loop (FlAsH 2) and the C-edge loops (FlAsH 4 and FlAsH 5) (**Figure 3a**). Interestingly, β-arrestins in lipid rafts, like in non-raft membrane domains, adopted similar conformations with different AT1R ligands, consistent with the agonist insensitivity of catalytically activated β-arrestins. The middle loop (FlAsH 2), C-edge loops (FlAsH 4 and FlAsH 5), and C-tail (FlAsH 6) displayed significant differences at raft and non-raft domains of the cell membrane, suggesting that these motifs might play a role in anchoring β-arrestins to the lipid bilayer (**Figure 3b-e**).

**Figure 3:**
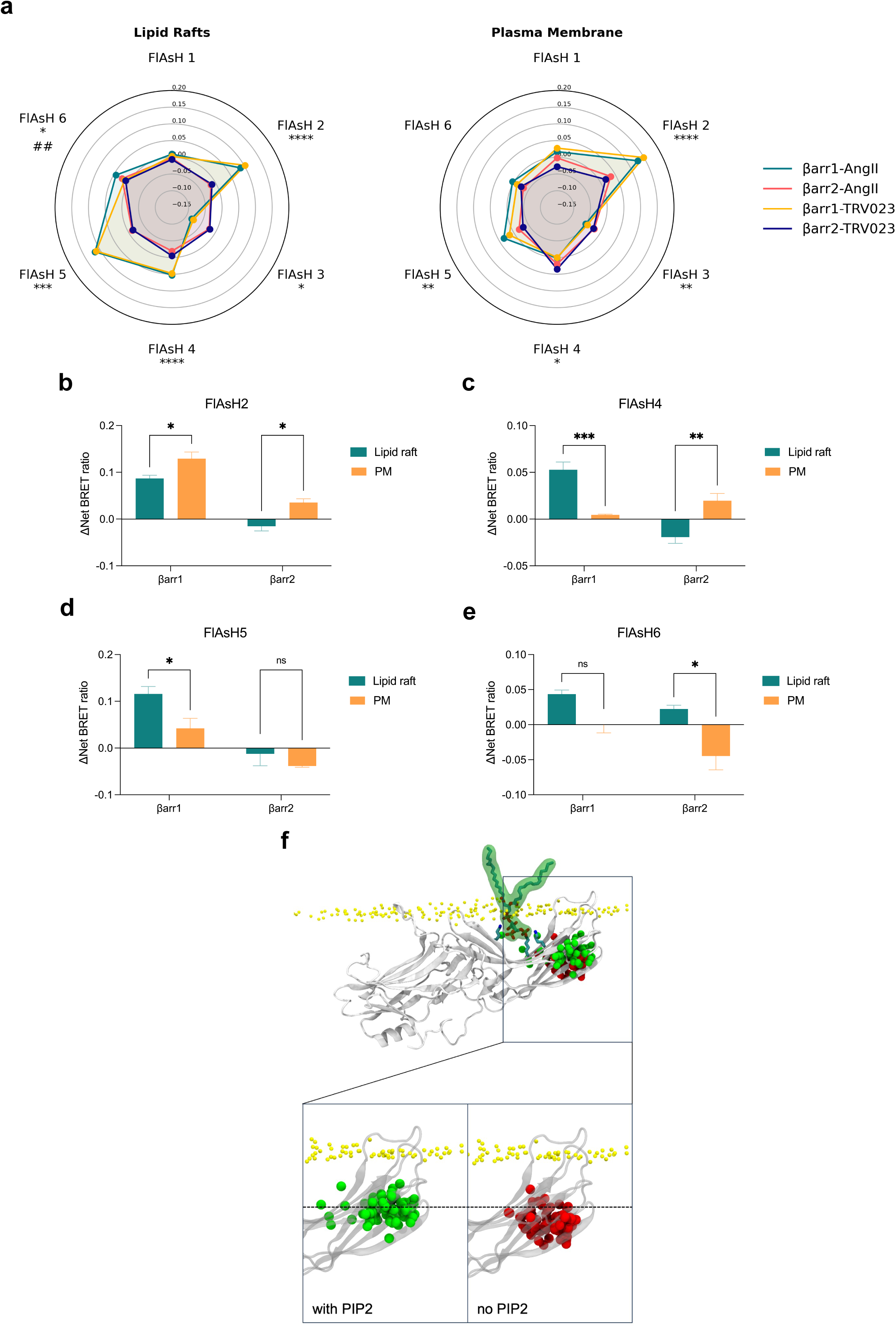
β-arrestin conformations are dependent on the lipid environment of the plasma membrane. **(a)** Radar plots from NanoBiT FlAsH assay demonstrate distinct conformations of β-arrestins 1 and 2 in non-raft membrane and lipid rafts after 2-10 min stimulation of AT1R with 1 μM AngII or 10 μM TRV023. No statistically significant agonist effect was observed in β-arrestin 1 and 2 conformations. Two-way ANOVA with Tukey’s multiple comparison test. * denotes the statistically significant differences between the β-arrestin isoforms. ^#^ denotes the statistically significant differences between the AT1R agonists. *P<0.05; **P<0.005; ***P<0.0005; ****P<0.0001; ^##^P<0.005. **(b-e)** Comparison of the BRET signals of FlAsH 2, FlAsH 4, FlAsH 5, and FlAsH 6 of β-arrestins 1 and 2 at lipid rafts and PM. Cells were stimulated with AngII. Data represents mean ± SEM, n=3 independent biological replicates for lipid rafts, n values for PM similar to figure 2. Two-way ANOVA with Šídák’s multiple comparison test to compare between PM vs lipid raft for each isoform. *P<0.05; **P<0.005; ***P<0.0005; ns, not significant. **(f)** MD simulations of β-arrestin 1 anchored to the PM (3 x 500ns) with (green) and without (red) PIP2. The simulation was aligned using membrane atoms. The position of the Cα atom of N223 (FlAsH 4) during the last half of each replicate is depicted in 10 ns intervals.

Since the most notable differences were observed in the C-edge loops that interact with the membrane, we hypothesized that the distinct lipid composition of non-raft membrane and lipid rafts influences their conformations. Phosphoinositides are heterogeneously distributed in the PM, with saturated lipids like phosphatidylinositol (3,4,5)-trisphosphate (PIP3) partitioned into cholesterol-rich, ordered lipid rafts, while unsaturated phospholipids such as phosphatidylinositol 4,5-bisphosphate (PIP2) localize in the disordered region of the PM^46,53,54^. To further explore this hypothesis, we performed MD simulations (3 x 500 ns) of β-arrestin 1 anchored to the membrane with and without PIP2 and monitored the position of the Cα atom of Asn223 (FlAsH 4) in the C-edge loops (**Figure 3f**). Interestingly, when comparing the position of the FlAsH 4 insertion across the simulation frames, we observe that the corresponding atom in simulations including PIP2 consistently assumes positions farther from the membrane. This suggests that PIP2 contributes to increased stabilization of the C-edge’s anchoring to the membrane. Given PIP2’s role as an additional anchoring point between the finger loop and C-edge, and its ability to act as a microswitch in β-arrestin activation, this finding aligns with its known functions^55–57^. Such results are further in line with FlAsH readouts, which demonstrate that changes in FlAsH signal induced by different lipid environments are strongest in the region of the C-edge (FlAsH 4 and FlAsH 5). These findings suggest that catalytically active β-arrestins exist in different lipid microdomains and that their unique lipid composition and membrane properties can affect β-arrestin conformations.

### ERK signaling at the plasma membrane is mediated by catalytically activated β-arrestins

To determine the consequences of location bias on AT1R signaling, we assessed the activation of ERK1/2 at various cellular locations. Using a previously published FRET-based ERK activity reporter (EKAR)^58^, we generated BRET-based ERK biosensors targeted to the cytosol, nucleus, early endosomes, and PM (**Figure 4a-d**). These biosensors consist of an N-terminal NLuc, followed by a proline-directed WW phospho-binding domain, a flexible 72-glycine linker, a substrate phosphorylation peptide from Cdc25C containing the ERK1/2 consensus targeting sequence (PDVPRTPVGK), the ERK-specific docking site (FQFP) and an mVenus BRET acceptor (**Extended Data Figure 6a**). This construct is naturally expressed in the nucleus due to the nuclear localization of the WW domain. A C-terminal location-targeting sequence was inserted to express the biosensor in the cytosol (nuclear export sequence, NES), endosomes (2xFYVE), or PM (CAAX). Following ERK1/2 activation, ERK1/2 phosphorylates the substrate peptide, which is then bound by the WW domain, resulting in a conformational change in the biosensor and subsequent increase in BRET efficiency. The localization of the EKAR biosensors to different cellular compartments was validated using confocal microscopy (**Extended Data Figure 6b-e**).

**Figure 4:**
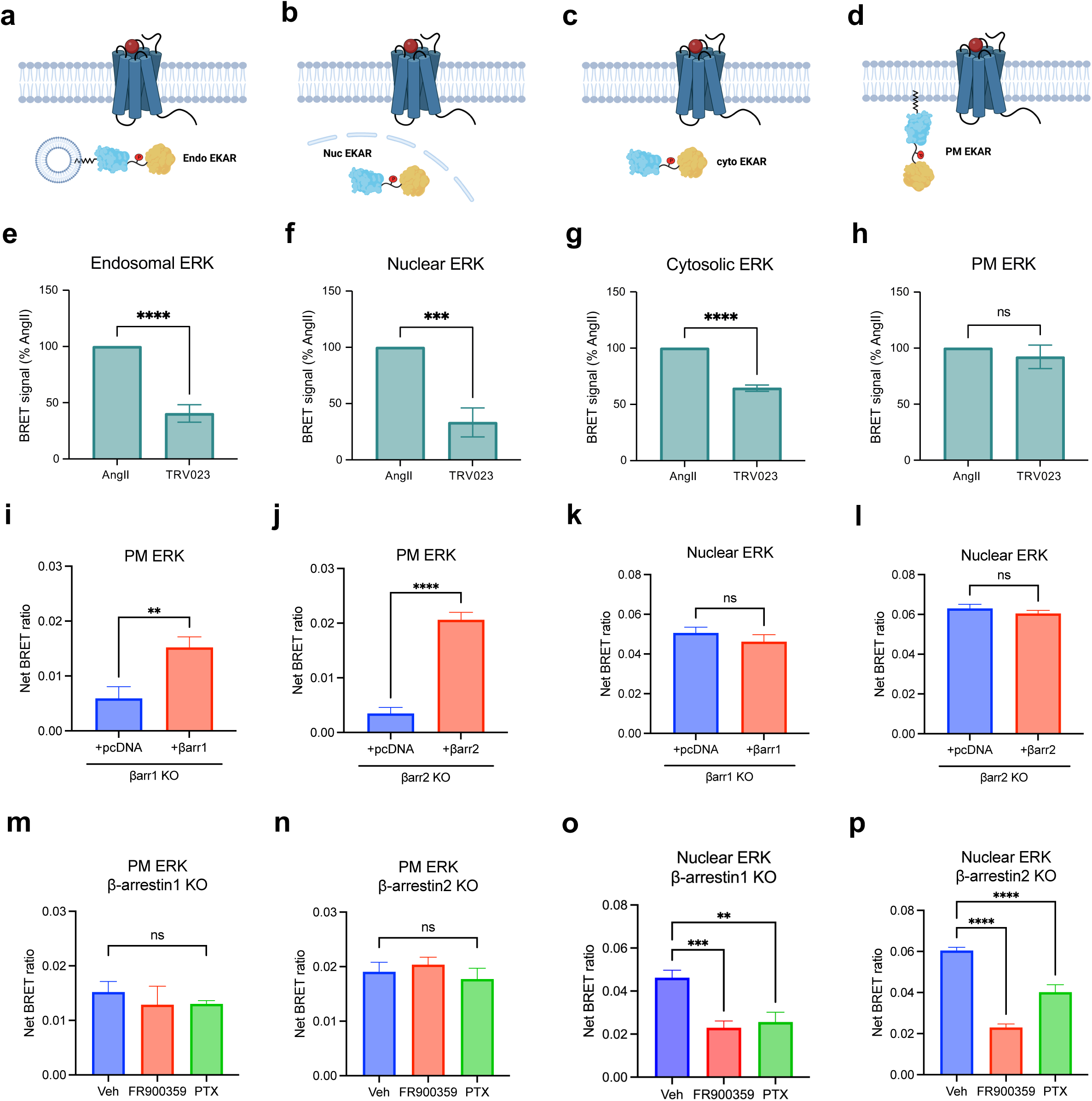
AngII and TRV023 promote distinct ERK signaling profiles at different cellular locations. (**a-d**) Diagrams of EKAR BRET biosensors subcellular targeted to the early endosomes, the nucleus, the cytosol, and the PM to measure location-specific ERK activity induced by AT1R ligands. **(e-h)** Area-under-the-curve (AUC) quantification of endosomal, nuclear, cytosolic, and PM ERK activity during the 50-minute stimulation of 1 μM AngII or 10 μM TRV023. Data was normalized to AngII as max signal and represents mean ± SEM of *n* independent biological replicates, n=4 for PM and cytosolic ERK, n=5 for nuclear and endosomal ERK. Unpaired Student’s t-tests comparing AngII versus TRV023. ***P<0.0005; ****P<0.0001; ns, not signficant. **(i-l)** PM and nuclear ERK activity measured in β-arrestin 1 KO or β-arrestin 2 KO HEK293 cells upon rescue with pcDNA control or FLAG-β-arrestin 1 or FLAG-β-arrestin 2. Cells were stimulated with 1 μM AngII for 5 min (PM ERK) or 30 min (nuclear ERK). Data represents mean ± SEM of *n* independent biological replicates, n=9 for PM ERK, n=7 for nuclear ERK. Unpaired two-tailed t-tests comparing pcDNA vs. β-arrestin rescue. **P<0.005; ****P<0.0001; ns, not significant. **(m-p)** Effect of Gq inhibition and Gi inhibition using FR900359 and PTX, respectively, on PM and nuclear ERK signaling in β-arrestin 1 KO or β-arrestin 2 KO HEK293 cells overexpressing FLAG-βarrestin 1 or FLAG-βarrestin 2. Cells were stimulated with 1 μM AngII for 5 min (PM ERK) or 30 min (nuclear ERK). Data represents mean ± SEM of *n* independent biological replicates, n=7 for nuclear EKAR, n=9 for PM ERK with vehicle, n=5 for PM ERK with the inhibitors. One-way ANOVA with Šídák’s posthoc test comparing inhibitors vs vehicle. ***P<0.0005; ****P<0.0001; ns, not significant.

We found that different AT1R agonists promoted distinct location-biased ERK signaling profiles. The β-arrestin-biased agonist TRV023 promoted significantly lower ERK activity in early endosomes, the nucleus, and cytosol compared to AngII, suggesting the role of G_q_ in mediating ERK activity at these locations (**Figure 4e-g**). While AngII promoted the most ERK activation in endosomes, TRV023 promoted the highest ERK activity at the PM, followed by the cytosol (**Extended Data Figure 6f, g**). Interestingly, both ligands induced similar levels of PM ERK activation, suggesting that ERK signaling at the PM could be primarily mediated by membrane-localized β-arrestins (**Figure 4h**). Utilizing HEK293 cells with either β-arrestin 1 or 2 knocked out (KO), we found that PM ERK activity was almost completely abolished in both cell lines and was rescued following overexpression of FLAG-β-arrestin 1 or FLAG-β-arrestin 2 (2.6-fold increase with β-arrestin 1, 3.6-fold increase with β-arrestin 2) (**Figure 4i, j, Extended Data Figure 7a, b**). Meanwhile, nuclear ERK activity was unaffected in the KO cells and was slightly decreased with β-arrestin overexpression (**Figure 4k, l, Extended Data Figure 7c**). Furthermore, inhibition of G proteins using the G_q_-selective inhibitor FR900359 or G_i_-selective pertussis toxin (PTX) had no measurable effect on the PM ERK activity rescued by overexpression of β-arrestin 1 or 2 (**Figure 5m, n**). On the contrary, G_q_ and Gi inhibition significantly impaired nuclear ERK signaling (**Figure 5o, p**). Endosomal and cytosolic ERK signaling also displayed a pattern similar to the observations with nuclear ERK although G_q_ played a more significant role than G_i_ for endosocmal ERK activation (**Extended Data Figure 7d-g**). Overall, these findings demonstrate that the AT1R ligands promote biased activation of ERK signaling at different cellular locations, with β-arrestin and G_q_ both promoting ERK activity at the endosomes, nucleus, and cytosol, while only β-arrestins are required to promote ERK signaling at the PM.

**Figure 5:**
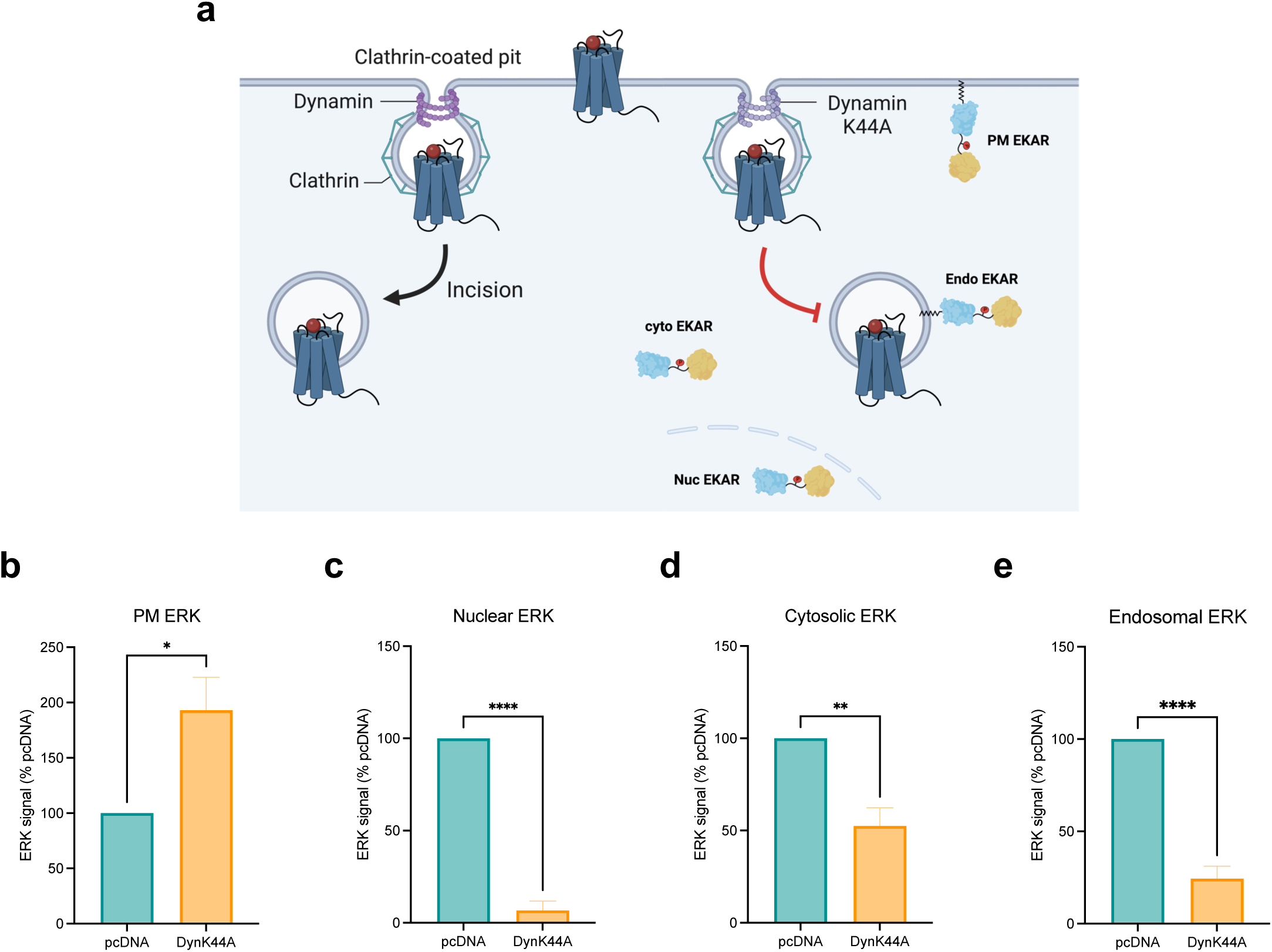
Endocytosis is essential for regulating ERK signaling in subcellular locations. (**a**) Schematic of EKAR BRET assay with or without endocytosis inhibition. Cells were transfected with FLAG-AT1R, EKAR biosensors, and dynamin K44A to inhibit receptor internalization or pcDNA 3.1 as control. **(b-e)** AUC quantification of ERK activity in early endosomes, nucleus, cytosol, and PM with endocytosis inhibition. Cells were stimulated with 1 μM AngII for 5 min (PM ERK) or 30 min (nuclear, cytosolic, endosomal ERK). Data was normalized to AngII-stimulated pcDNA condition as 100% and is shown as mean ± SEM of *n* independent biological replicates, n=4 for PM and cytosolic ERK, n=5 for nuclear and endosomal ERK. Statistical analysis was performed using unpaired two-tailed t-tests to compare pcDNA vs. dynamin K44A. *P<0.05; **P<0.005; ****P<0.0001.

### Subcellular pools of ERK signaling are differentially regulated by endocytosis

We finally investigated the role of endocytosis as a mechanism for regulating and distributing the different pools of ERK signaling in subcellular locations. Inhibition of endocytosis was achieved by overexpressing Dynamin K44A (DynK44A), a dominant-negative mutant of the GTPase Dynamin, which facilitates the scission of clathrin-coated pits and release into the PM^59^ (**Figure 5a**). Endocytosis inhibition by Dynamin K44A substantially increased PM ERK activity (**Figure 5b**). On the other hand, impairment of endocytosis abrogated nearly all endosomal and nuclear ERK activity and partially reduced ERK activation in the cytosol (**Figure 5c-e)**. These data reveal that agonist-induced endocytosis is crucial for endosomal and nuclear ERK activation and plays a contributing role, though not essential, for cytosolic ERK activation. In contrast, inhibition of endocytosis increased PM ERK activity, likely due to decreased sequestration of ERK at the PM.

## DISCUSSION

In the classical view of GPCR signal transduction, signaling cascades are initiated at the plasma membrane (PM) through interactions between the receptor, G proteins, β-arrestins, and GPCR kinases (GRKs). However, emerging research has demonstrated that GPCR signaling is not confined to the PM; it can extend into various subcellular compartments such as endosomes and the nucleus, adding layers of complexity to cellular signaling networks^9–11,60,61^. This study provides concrete evidence that β-arrestins play a pivotal role in this location bias by adopting distinct trafficking patterns and conformations that drive functionally selective downstream ERK signaling responses (**Figure 6**).

**Figure 6:**
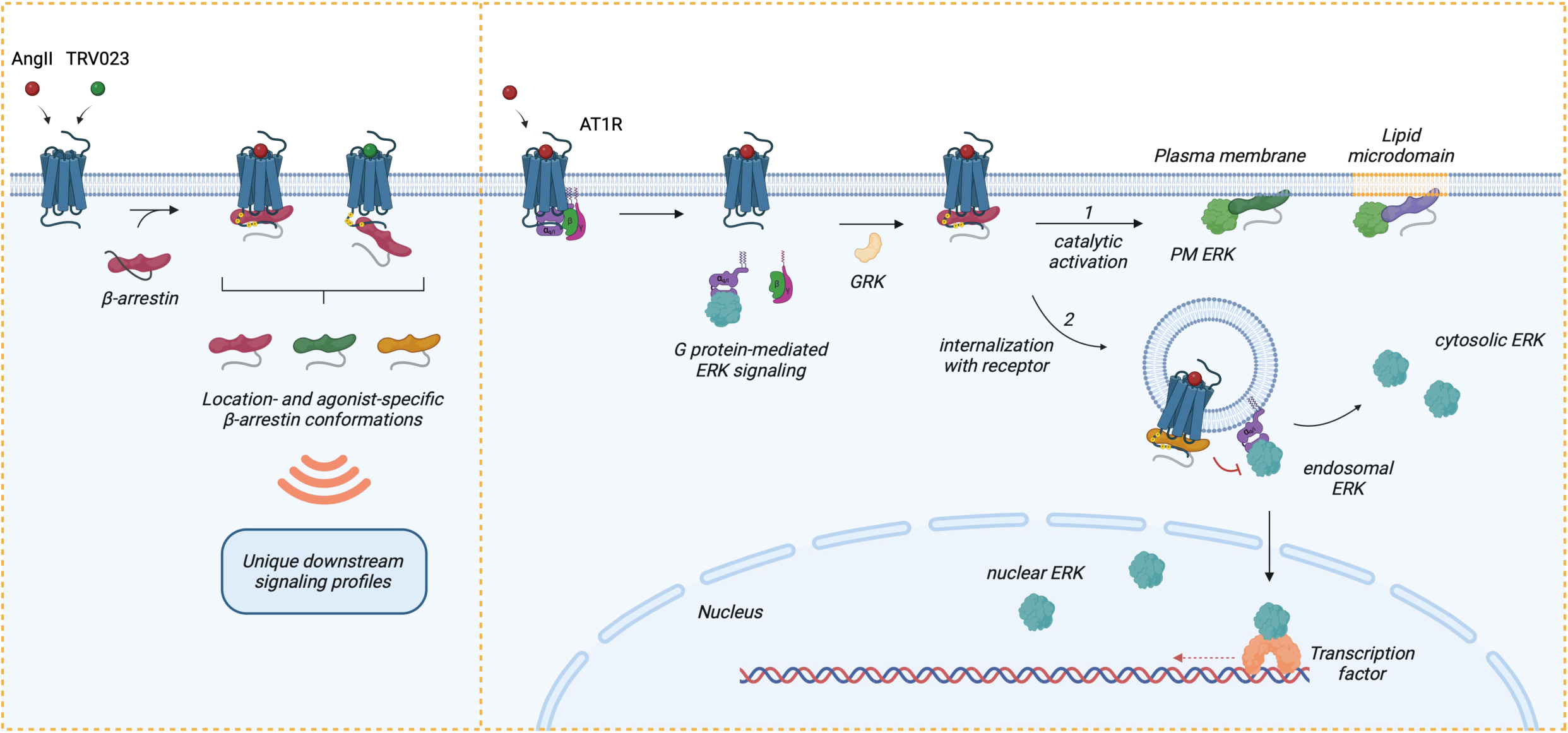
Model of regulation of location-biased ERK activation downstream of AT1R. Upon stimulation with agonists, AT1R activates Gα_q_ and Gα_i_, resulting in the dissociation of the G protein subunits and promoting endosomal ERK signaling (after receptor internalization mediated by β-arrestins), which can propagate into the cytosol and nucleus. Following G protein activation, GRKs phosphorylate the receptor C-tail, which promotes the recruitment of β-arrestins. Depending on the agonist, β-arrestins can form two main complexes with the receptor: a core conformation induced by AngII and a hanging tail conformation induced by TRV023. Each configuration stabilizes distinct conformational profiles of receptor-bound β-arrestins. Subsequently, β-arrestins either (1) undergo catalytic activation by the receptor, allowing them to translocate across the PM and lipid microdomains independently of the receptor, or (2) co-traffic with the internalized receptor into endosomes. Depending on the membrane lipid environment and subcellular locations, catalytically active β-arrestins and receptor-associated β-arrestins adopt unique conformational signatures that dictate their function, such as regulating the ERK/MAPK signaling pathway. While ERK activity at the PM is mainly promoted by catalytically active β-arrestins, endosomal and nuclear ERK signaling is activated by Gα_q_ and Gα_i_ and can be inhibited by β-arrestins. Receptor endocytosis distributes subcellular pools of ERK signaling from the PM into the endosome, cytosol, and nucleus. In summary, β-arrestin plays critical roles in regulating the intensity, duration, and location bias of AT1R signaling through desensitization, internalization, and catalytic activation initiating its own pattern of ERK signaling, thus modulating the cellular response.

Our findings reveal the existence of separate pools of catalytically active β-arrestins at the PM, distinct from those associated with receptor trafficking to endosomes, which is consistent with previous studies^19,43^. These catalytically active β-arrestins, activated by GPCRs following transient receptor engagement, exhibit unique conformational states compared to β-arrestins that co-traffic with the receptor. This observation is supported by our NanoBiT FlAsH experiments, which highlight the differential conformations of membrane-bound β-arrestins that are independent of ligand bias, while receptor– and endosome-associated β-arrestins display ligand bias-dependent effects, underscoring the importance of receptor interaction in determining β-arrestin conformational changes. This is consistent with research that have shown the propagation of ligand-specific conformational rearrangements in β-arrestins through receptor interaction^62,63^.

Further structural insights, derived from molecular dynamics (MD) simulations, support a model where key structural elements of β-arrestin 1—including the finger loop, C-loop, and C-edge loops— insert into the lipid bilayer, stabilizing its catalytically active form at the PM, which is consistent recent structural and single-molecule microscopy studies^43,64^. This membrane insertion shifts the middle loop toward the N-terminal region, resulting in a robust increase in BRET signal, a finding corroborated by both our experimental and computational data. Interestingly, different agonists appear to stabilize unique receptor-β-arrestin complexes, with AngII promoting a core conformation that fully engages the receptor transmembrane core, whereas TRV023 stabilizes a low-affinity hanging tail complex. This is consistent with previous findings that AngII and G-protein-biased agonists promote open conformations of the AT1R transmembrane core with an accessible transducer binding site while β-arrestin-biased agonists such as TRV023 stabilize occluded conformational states of the receptor core which show lower β-arrestin binding efficacy^21^. Future research should explore a broader panel of G protein– and β-arrestin-biased agonists to further delineate these interactions and investigate whether specific receptor-β-arrestin conformations preferentially drive catalytic activation versus co-trafficking and whether this depends on agonist-induced phosphorylation patterns of the receptor C-tail (the barcode hypothesis)^32,65^.

Another key discovery is the presence of catalytically active β-arrestins in the PM, displaying distinct conformations in lipid raft and non-raft membranes. Structural changes compared to the basal conformation β-arrestin in the cell are particularly notable in regions such as the finger loop, C-edge loops, and C-terminal tail. Our data suggest that the conformational insensitivity of β-arrestins to AT1R ligand bias observed at non-raft regions is also present in lipid rafts, where membrane properties and lipid composition—especially the presence of PIP2—play crucial roles in anchoring these regions. Although the structure of β-arrestin’s C-terminal tail remains unresolved due to its disordered nature, its distinctive orientation in lipid rafts suggests potential interactions with membrane phospholipids.

Biased AT1R agonists promote distinct patterns of ERK activity across cellular compartments. While Gq and β-arrestins both modulate ERK signaling in the cytosol, endosomes, and nucleus, only β-arrestins, likely those in the catalytically active pool, drive membrane-localized ERK signaling. This observation aligns with the finding that receptor-independent recruitment of β-arrestin 2 to the PM is sufficient to activate ERK, and that PIP2 can stabilize an active β-arrestin conformation and influence the stability of the GPCR-β-arrestin complex^55,66^. Previous studies have established that endocytosis is a key driver of biased AT1R signaling, mediating differences in β-arrestin binding efficacy among various agonists^67^. Our work expands on this by demonstrating that agonist-stimulated endocytosis is also a critical factor in propagating spatially biased AT1R signaling, effectively distributing different ERK signaling pools from the PM into endosomes, the nucleus, and the cytosol.

Overall, our study underscores the intricate dynamics between β-arrestin isoforms, ligand bias, and location bias in fine-tuning the signaling responses downstream of GPCRs. The AT1R and AngII-induced ERK signaling cascades we studied are known to drive physiological and pathological processes, including muscle cell hypertrophy and adverse cardiac remodeling, with significant clinical relevance in conditions such as hypertension, diabetes, heart failure, and atherosclerosis^68,69^. The increasing focus on developing biased agonists, particularly β-arrestin-biased ligands, reflects a growing interest in designing pharmaceuticals that selectively target therapeutic pathways while minimizing signaling through other pathways^70^. The divergence between β-arrestin isoforms offers additional opportunities for ligand specificity, as β-arrestin 1 and β-arrestin 2 exhibit distinct functions and conformations at the AT1R and other GPCRs^71,72^. Additionally, the concept of location bias opens new avenues in drug discovery, with efforts targeting specific subcellular pools of GPCR signaling— such as β1AR, NK1R, and μ-OR—aiming to improve therapeutic efficacy and specificity^4^.

## ACKNOWLEDGEMENTS

We thank Dr. Robert Lefkowitz for the guidance and thoughtful feedback throughout this work and for providing the anti-β-arrestin antibody (A1CT); Dr. Louis Luttrell for sharing the Rluc-β-arrestin2-FlAsH plasmids; Dr. Howard Rockman for providing the ARBB1 KO and ARBB2 KO HEK293 cells; Dr. Xinyu Xiong for cloning the SmBiT-β-arrestin2-FlAsH plasmids; Claudia Lee for the reagents and thoughtful feedback throughout this work; and Nour Nazo for laboratory assistance. This work was supported by a National Institute of Health grant 1R01GM122798 and Mandel Scholar Award (S.R.), the American Heart Association Predoctoral Fellowship 23PRE1019796 (U.P.), the Mandel Foundation from Duke Cardiovascular Research Center (S.R.), Sara Borrell grant CD22/00007 funded by the Institute of Health Carlos III (ISCIII), and grant 2021 SGR 00046 funded by Agència de Gestió d’Ajuts Universitaris i de Recerca Generalitat de Catalunya, Instituto de Salud Carlos III FEDER (PI18/00094). Figures were created using GraphPad Prism 10, BioRender, Python, and Visual Molecular Dynamics (VMD).

## AUTHOR CONTRIBUTIONS

Conceptualization, U.P., S.R.; Methodology, U.P., S.R.; Investigation, U.P., A.C., T.M.S., D.S.E., S.D., S.G., C.H., J.G.; Writing — Original Draft, U.P.; Writing – Reviewing & Editing, U.P., A.C., T.M.S., D.S.E., S.D., S.G., C.H., J.G., S.R.; Visualization, U.P., A.C., T.M.S; Supervision and Funding Acquisition, S.R.

## DECLARATION OF INTERESTS

The authors declare no competing interests.

## MATERIALS AND METHODS

### Plasmid Constructs

#### Generation of SmBiT-βarr-FlAsH constructs

Rluc-β-arrestin2-FlAsH constructs (FlAsH1-6)^27^ were provided by Dr. Louis Luttrell (Medical University of South Carolina, Charleston, SC). SmBiT-β-arrestin2-FlAsH (rat) constructs were cloned by removing the N-terminal RLuc of FlAsH1-6 and replacing it with SmBiT using restriction digest. To generate SmBiT-β-arrestin 1-FlAsH (rat) constructs, the CCPGCC motif was inserted after amino acids G39, K138, K170, N223, T261, and G409 of SmBiT-β-arrestin 1 plasmid using overlap extension PCR.

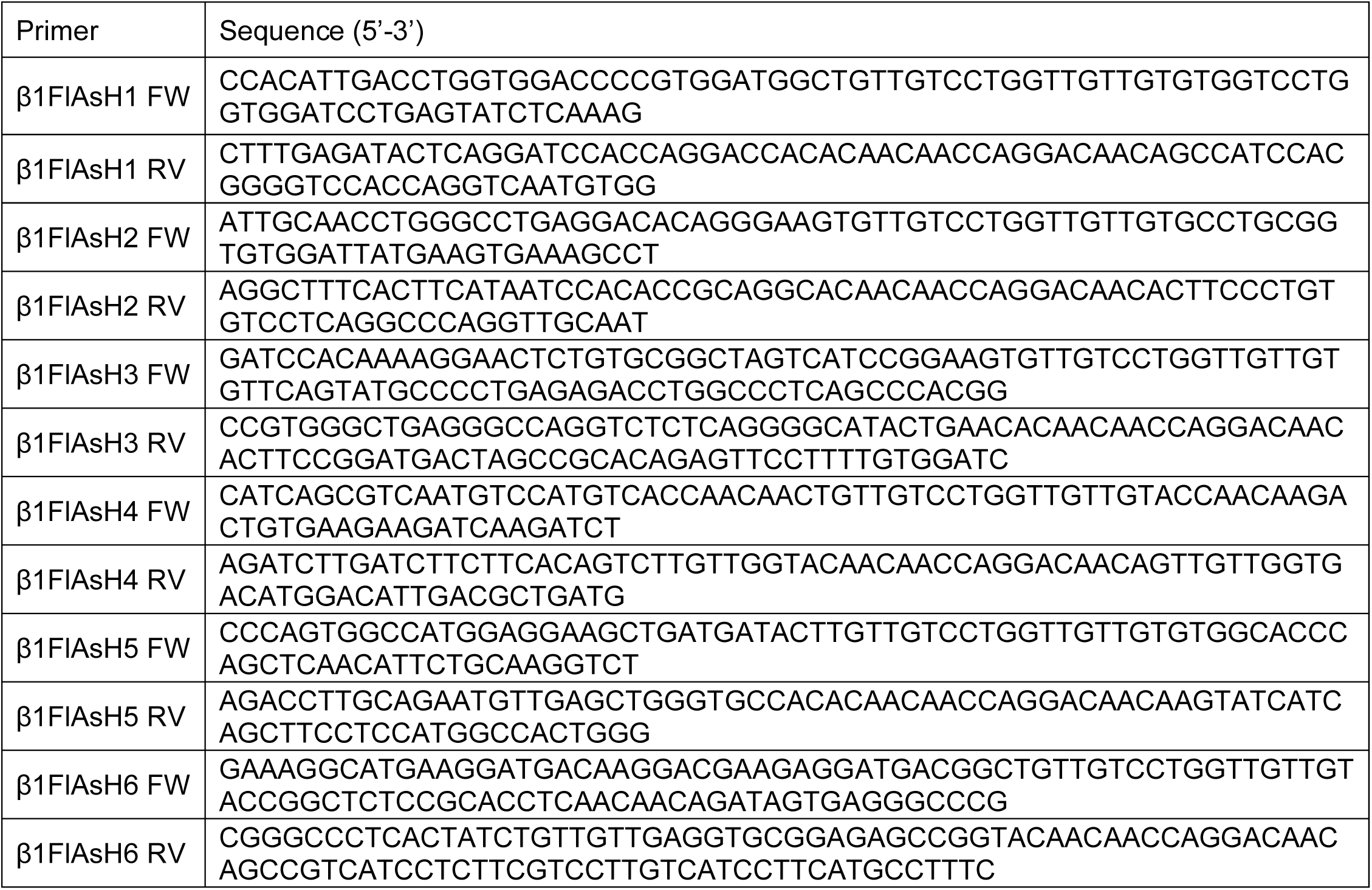

#### Generation of EKAR BRET biosensors

Previously published EKAR FRET biosensors for cytosolic and nuclear ERK1/2 activity^58^ were used to clone their BRET versions by removing the N-terminal mCerulean through restriction digest and inserting an NLuc. EKAR-CAAX and EKAR-2xFYVE constructs were cloned using overlap extension PCR. In short, the NLuc sequence was amplified from the cyto EKAR BRET plasmid. Then the ECFP was removed from pm-EKAR4 and endo-EKAR4 plasmids previously published from Dr. Jin Zhang’s laboratory^11^ and replaced with NLuc using PCR.

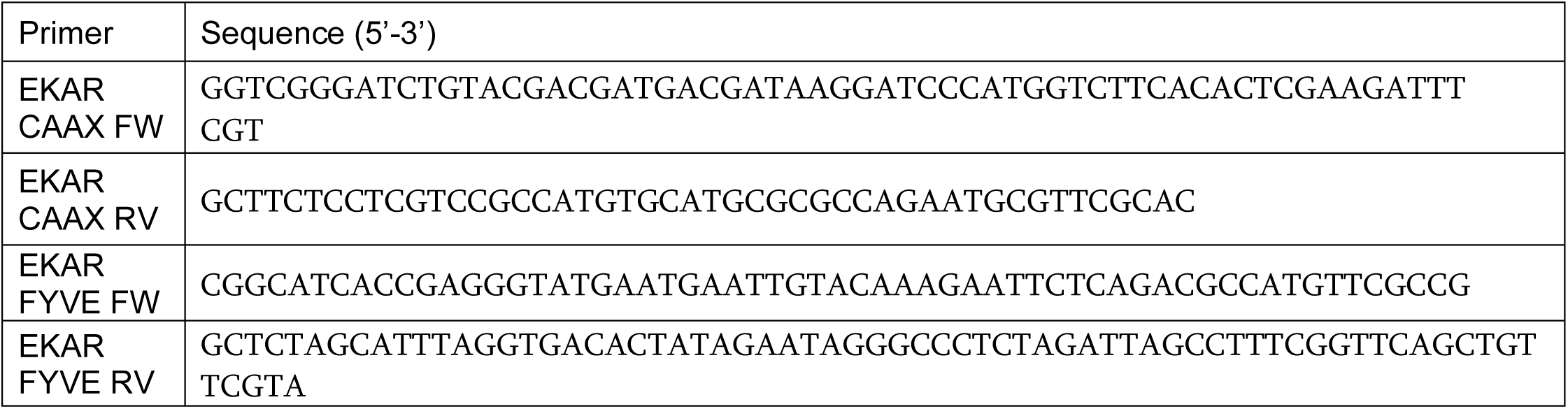

#### Generation of β-arrestin ΔFL mutants

Overlap extension PCR was used to delete the finger loop sequence from SmBiT-β-arrestin 1 (rat) or SmBiT-β-arrestin2 (mouse). This protocol utilized 4 chimeric primers designed to amplify two DNA fragments directly before or after the finger loop sequence of each SmBiT-β-arrestin plasmid and introduce an overlapping sequence into the target fragments. In the second step, a ligation PCR was performed, which utilized the overlapping sequences as the primers to facilitate the extension of the PCR and join the two adjacent fragments together.

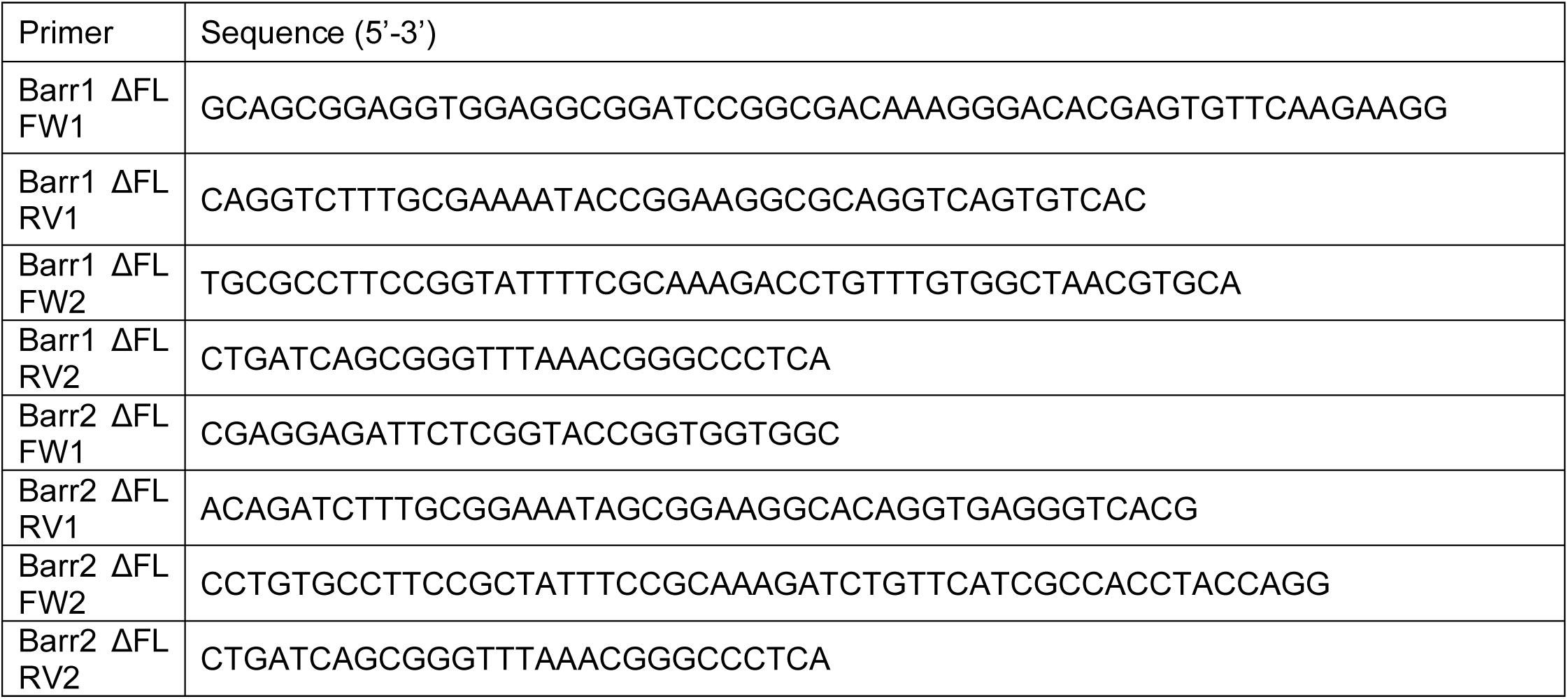

### AT1R Ligands and Inhibitors

Angiotensin II was purchased from Sigma-Aldrich and TRV120023 was synthesized by GenScript. Both ligands were reconstituted in ultrapure water, and aliquots were stored at –20°C. FR900359 was purchased from Cayman Chemical. Pertussis toxin was purchased from List Biological Laboratories.

### Cell Culture and Transfection

ARRB1 KO HEK293 and ARRB2 KO HEK293 cells were provided by Dr. Howard Rockman’s laboratory. Human Embryonic Kidney cells (WT HEK293T, ARRB1 KO HEK293, and ARRB2 KO HEK293) were grown in minimum essential media (MEM) supplemented with 10% fetal bovine serum (FBS) and 1% penicillin/streptomycin (P/S) at 37°C and 5% CO_2_. Transient transfections were performed using polyethylenimine (PEI). Briefly, MEM was replaced 30 minutes prior to transfection. Plasmid constructs were suspended in Opti-MEM (Gibco) to a final volume of 100 μL. In a separate tube, 100 μL of PEI in Opti-MEM was prepared at a PEI:DNA ratio of 3:1. After 5 minutes, PEI solution was added to the plasmid DNA, gently mixed, and allowed to incubate at room temperature (RT) for 30 minutes. The PEI and DNA mixtures were then added to cells with gentle swirling.

### NanoBiT β-arrestin Recruitment Assays

HEK293T cells seeded in 6-well plates were transiently transfected with FLAG-AT1R, either SmBiT-β-arrestin 1 or SmBiT-β-arrestin 2 and a location marker (LgBiT-CAAX or 2xFYVE-LgBiT). For β-arrestin recruitment to the receptor, cells were transfected with AT1R-LgBiT and either SmBiT-β-arrestin 1 or SmBiT-β-arrestin 2. 24 hours after transfection, cells were plated onto clear bottom, white-walled, 96-well plates (Costar) at 100,000 cells/well in clear MEM supplemented with 2% FBS, 1% penicillin/streptomycin, 10 mM HEPES, 1x GlutaMax, and 1x Antibiotic-Antimycotic (Gibco). The following day, the media were aspirated, and cells were incubated at RT with 80 μL of coelenterazine h (2.5 μM final concentration) in Hanks’ balanced salt solution (HBSS) (Gibco) supplemented with 20 mM HEPES for 5 minutes. Luminescence signals were read using a BioTek Synergy Neo2 plate reader at 37°C. Three prereads were taken to quantify the baseline luminescence before adding 20 μL of ligands at the appropriate concentrations. Baseline luminescence was divided from each read following ligand addition to calculate a change in luminescence over baseline and then normalized to vehicle treatment.

### NanoBiT FlAsH BRET Assays

HEK293T cells were seeded in 6-well plates and were transiently transfected 24 hours later using PEI. For experiments at the PM, early endosomes and lipid rafts, cells were transfected with FLAG-AT1R, either SmBiT-β-arrestin 1-FlAsH probes or SmBiT-β-arrestin2-FlAsH probes and a location marker (LgBiT-CAAX, 2xFYVE-Lgbit or MyrPalm-LgBiT). For FlAsH experiments at the receptor, cells were transfected with AT1R-LgBiT and either SmBiT-β-arrestin 1-FlAsH or SmBiT-β-arrestin 2-FlAsH. 24 hours after transfection, cells were plated onto clear-bottomed, rat tail collagen-coated, white-walled, 96-well plates (Costar) at 100,000 cells/well in MEM (Gibco) supplemented with 10% FBS and 1% P/S. The following day, cells were washed with 60 μL of HBSS (+ 20 mM HEPES, calcium, magnesium). After 60 μL of 2.5 μM FlAsH-EDT2 in HBSS was added for arsenical labeling or 60 μL of HBSS only was added for mock labeling, cells were incubated in a 37 °C, 5% CO_2_ incubator for 30 minutes. FlAsH-EDT2 was aspirated, and the cells were washed with 10-minute incubation of 120 μL of 250 μM 2,3 dimercapto-1-propanol (BAL) wash buffer (Sigma-Aldrich). For assay reading, cells were incubated at room temperature with 80 μL of coelenterazine h in HBSS (2.5 μM final concentration) for 5 minutes. Three prereads were then taken to measure the baseline signal. 20 μL of ligands were then added to a final concentration of 1 μM AngII and 10 μM TRV023. BRET signals were measured with a BioTek Synergy Neo2 at 37°C using a 480 nm wavelength filter (NLuc) and 530 nm wavelength filter (FlAsH-EDT2). BRET ratios were calculated by dividing the 530 nm signal by the 480 nm signal. Net BRET values were calculated by subtracting the vehicle BRET ratio from the ligand-stimulated BRET ratio. ΔNet BRET ratios were calculated by subtracting the FlAsH-labeled net BRET signals from the mock-labeled signals. For radar plot, an average of ΔNet BRET ratio was calculated at different time point post-stimulation: 2-10 minute for AT1R and PM, 30-40 minute for endosome.

### EKAR BRET Assays

HEK293T cells seeded in 6-well plates were transiently transfected with FLAG-AT1R, EKAR BRET biosensors tagged to different cellular locations, and dynamin K44A or pcDNA control. 24 hours later, cells were plated onto clear bottom, white-walled, 96-well plates (Costar) at 100,000 cells/well in clear MEM supplemented with 0.5% FBS and 1% P/S. The following day, the media were aspirated, and cells were incubated at RT with 80 μL of coelenterazine h (2.5 μM final concentration) in HBSS supplemented with 20 mM HEPES for 5 minutes. BRET signals were measured with a BioTek Synergy Neo2 plate reader at 37°C using a 480nm wavelength filter (NLuc) and 530nm wavelength filter (mVenus). Three prereads were taken to quantify the baseline BRET signals before 20 μL of ligands were added (1 μM AngII or 10 μM TRV023). BRET ratios were calculated by dividing the 530 nm signal by the 480 nm signal. Net BRET values were calculated by subtracting the vehicle BRET ratio from the ligand-stimulated BRET ratio. β-arrestin 1 or β-arrestin 2 KO HEK293 cells were seeded in 6-well plates and, after 24 hours, were transfected with FLAG-AT1R and EKAR biosensors. 24 hours later, cells were plated onto clear bottom, white-walled, collagen-coated 96-well plates (Costar) at 100,000 cells/well in clear MEM supplemented with 0.5% FBS and 1% P/S, with or without 200 ng/mL PTX. The next day, the media was removed and 20 uL of HBSS or FR900359 (1 μM final concentration) was added. Cells were then incubated at RT with 60 μL of coelenterazine h (2.5 μM final concentration) in HBSS for 5 minutes. Three prereads were taken, followed by the addition of 20 uL of AngII (1 μM final concentration). BRET signals were measured with a BioTek Synergy Neo2 plate reader, as previously described.

### Confocal Microscopy

35,000 HEK293T cells were plated on 35-mm glass-bottomed dishes coated with rat tail collagen (Sigma-Aldrich). For β-arrestin recruitment experiments, cells were transiently transfected 24 hours later with β-arrestin 1 RFP or β-arrestin 2 RFP and rGFP CAAX or rGFP 2xFYVE using PEI protocol. Cells were stimulated with AngII for 5 min (CAAX) or 45 min (2xFYVE). For visualization of EKAR BRET sensors, cells were transfected with 50 ng of each EKAR BRET biosensor using PEI. Forty-eight hours following transfection, the cells were washed once with PBS and serum starved for one hour. The cells were imaged with a Zeiss CSU-X1 spinning disk confocal microscope using the corresponding lasers to excite GFP (480nm) and RFP (561nm). Images were edited and analyzed using ImageJ (NIH, Bethesda, MD).

### Western Blotting

β-arrestin 1 KO and β-arrestin 2 KO HEK293 cells seeded in 6-well plates were transiently transfected with pcDNA 3.1, FLAG-β-arrestin 1 (β-arrestin 1 KO cells), or FLAG-β-arrestin 2 (β-arrestin 2 KO cells) using the PEI transfection method. After 48 hours, cells were washed with ice cold PBS and lysed in ice cold RIPA buffer supplemented with cOmplete EDTA-free protease inhibitors (Roche). The samples were rotated at 4° C for 1 hour and cleared of insoluble debris by centrifugation at 17,000g at 4° C for 15 minutes, after which the supernatant was collected. Protein was resolved on SDS-10% polyacrylamide gels, transferred to nitrocellulose membranes, and immunoblotted with the indicated primary antibody overnight at 4°C. Rabbit polyclonal anti-β-arrestin antibody (A1CT)^18^ (1:3000) and mouse monoclonal anti-α-Tubulin antibody (Sigma-Aldrich) (1:18000) were used for immunoblotting. Horseradish peroxidase-conjugated polyclonal mouse anti-rabbit-IgG or anti-mouse-IgG (Rockland) were used as secondary antibodies (1:3000). The nitrocellulose membranes were imaged by SuperSignal™ West Pico Plus chemiluminescent substrate (Thermo Fisher) using a ChemiDoc MP Imaging System (Bio-Rad). Following detection of β-arrestin signal, nitrocellulose membranes were stripped and reblotted for α-Tubulin.

### Molecular Dynamics Simulations

We modeled the complex of β-arrestin 1 and the membrane using a previously established approach^43^. We utilized the structure of β-arrestin 1 in complex with the neurotensin receptor 1 (PDB code: 6UP7)^40^. To facilitate interactions between β-arrestin 1 and the membrane, we utilized conformations of the C-edge loops and the finger loop from a previous equilibrated arrestin/membrane complex^43^. The sequence of β-arrestin 1 was modified to match the isoform used in the FlAsH in vitro experiments [UniProt: P29066]. The complexes were solvated (TIP3P water) and neutralized using a 0.15 M concentration of NaCl ions. Parameters for simulations were obtained from the Charmm36M forcefield^73^. We used a membrane consisting of 10% cholesterol, 38% palmitoyl-oleoyl-phosphatidylcholine, 28% dioleoyl-phosphatidylcholine, and 24% dioleoyl-phosphatidylethanolamine. Additionally, we included a PIP2 molecule in the lower and upper leaflet of the membrane, and the PIP2 molecule was placed based on previously established coorindates^40^. Simulations were run using the ACEMD3 engine^74^. All systems underwent a 100 ns equilibration in conditions of constant pressure (NPT ensemble, pressure maintained with Berendsen barostat, 1.01325 bar), using a timestep of 2 fs. During this stage, mobility restraints were applied to the backbone. This was followed with 3 × 500 ns of simulation for each system in conditions of constant volume (NVT ensemble) using a timestep of 4 fs. For every simulation we used a temperature of 310K, maintained using the Langevin thermostat. Hydrogen bonds were restrained using the RATTLE algorithm. Non-bonded interactions were cut off at a distance of 9 Ȧ, with a smooth switching function applied at 7.5 Ȧ.

### Statistical Analyses

Statistical analyses were performed using GraphPad Prism 10 (GraphPad Software). Values are reported as mean ± SEM. For split luciferase assays, dose-response curves were fitted to a log agonist versus stimulus with three parameters (span, baseline, and EC50) and baseline-corrected to zero. For NanoBiT FlAsH assays, statistical tests were performed using two-way ANOVA followed by Tukey’s multiple comparison test, one-way ANOVA followed by Tukey’s multiple comparison test, or unpaired two-tailed t-tests. For EKAR BRET assays, pairwise comparisons were performed using Student’s t-tests. Comparisons with two or more groups were performed using one-way ANOVA with Šídák’s multiple comparison test when comparing specific conditions. P<0.05 was considered to be statistically significant. Further details of statistical analysis and replicates are reported in the figure legends.

### Data Availability Statement

Source data are provided with this paper. The results of the MD simulations have been deposited at GPCRmd. The accession number is https://www.gpcrmd.org/dynadb/publications/1526/.

## REAGENTS AND RESOURCES

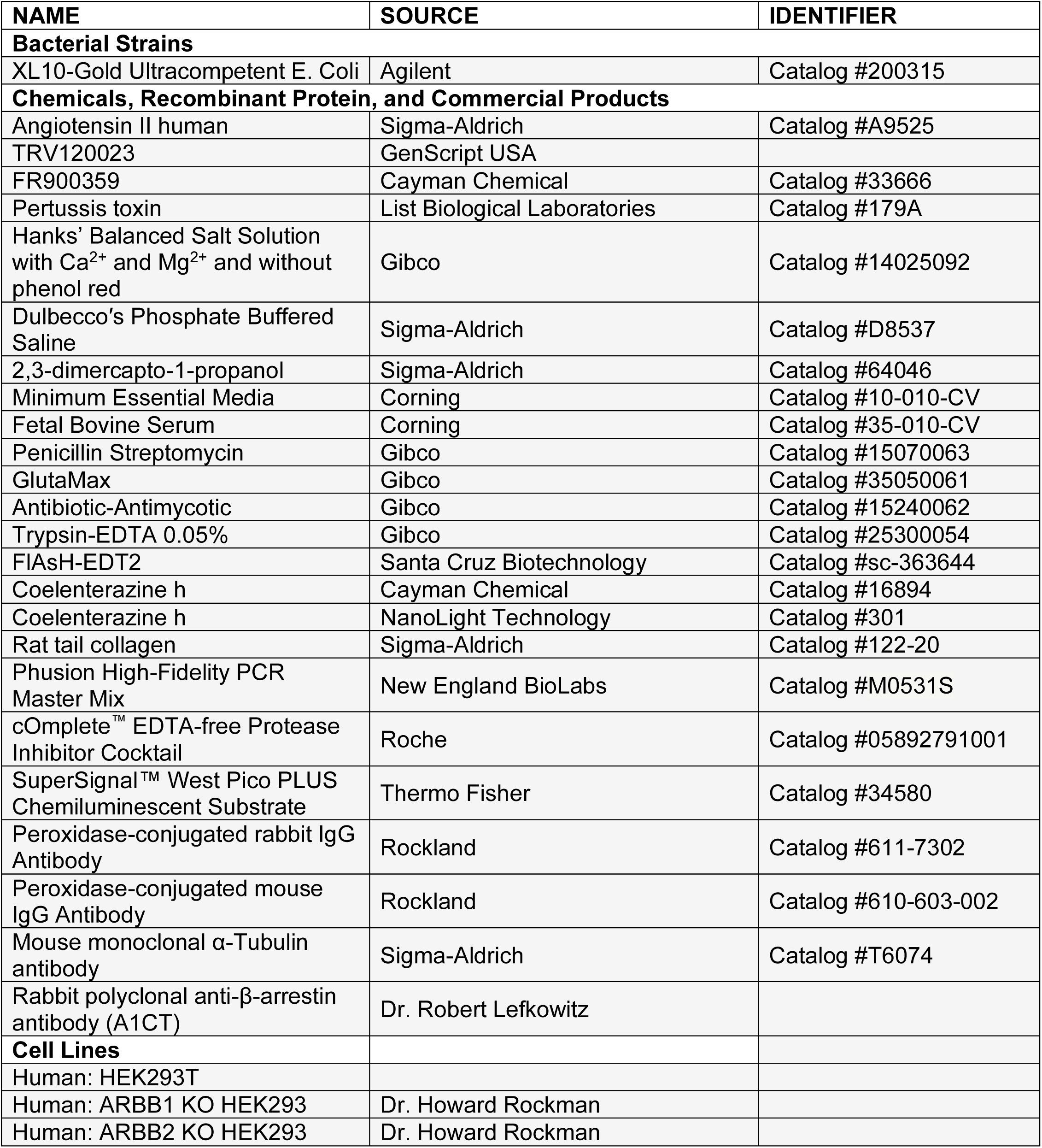

## SOFTWARE AND ALGORITHMS

| NAME | SOURCE | IDENTIFIER |
| --- | --- | --- |
| GraphPad Prism | GraphPad Software | https://www.graphpad.com/scientific-software/prism/ |
| ImageJ | <sup>75</sup> | https://imagej.nih.gov/ij/ |
| Adobe Illustrator | Adobe | https://www.adobe.com/ |
| Excel | Microsoft | https://www.microsoft.com/en-us/microsoft-365/excel |
| ImageLab | Bio-Rad | https://www.bio-rad.com/en-us/product/image-lab-software |
| BioRender | BioRender | https://biorender.com/ |
| Visual Molecular Dynamics | <sup>76</sup> | https://www.ks.uiuc.edu/Research/vmd/ |

## EXTENDED DATA FIGURE LEGENDS

**Extended Data Figure 1:**
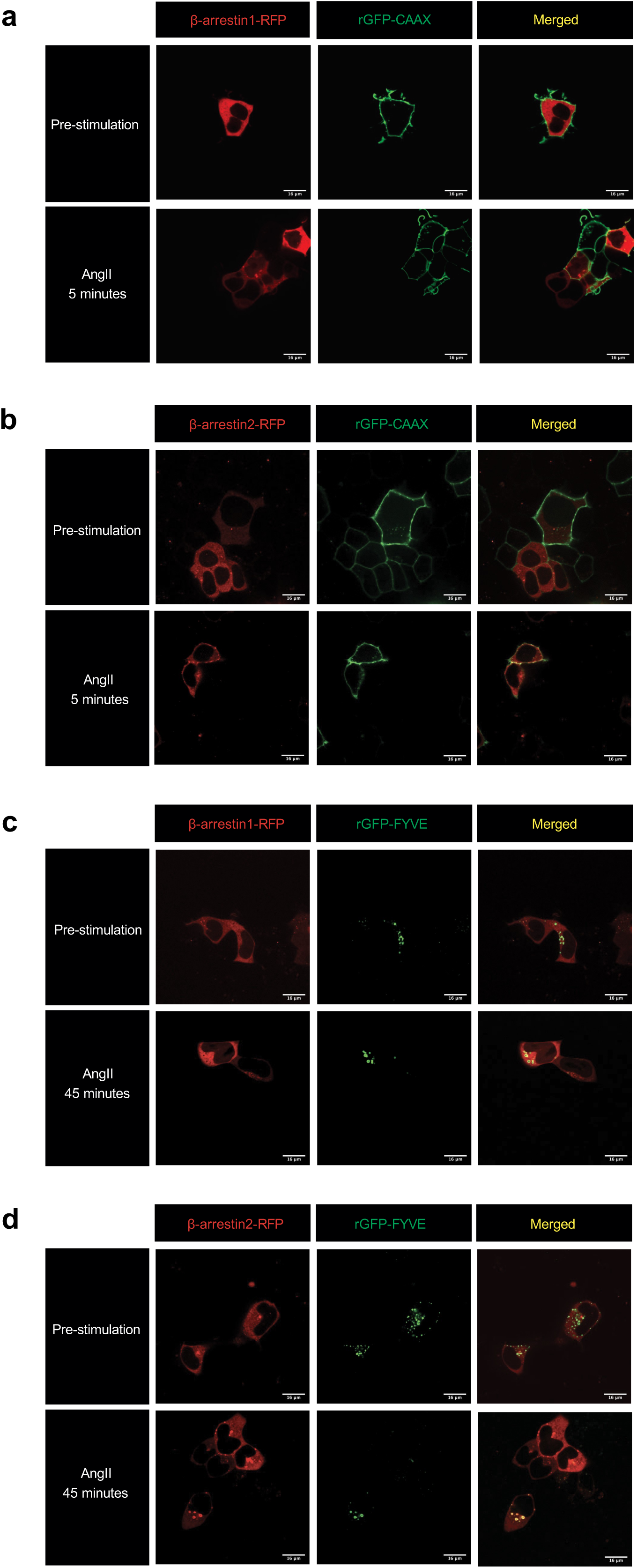
Confocal microscopy images of β-arrestin 1 and β-arrestin 2 trafficking to the plasma membrane and early endosomes. HEK293T cells transfected with FLAG-AT1R, PM marker rGFP-CAAX, and β-arrestin 1-RFP **(a)** or β-arrestin 2-RFP **(b)** pre-stimulation and after 5-minute stimulation of 1 μM AngII. HEK293T cells transfected with FLAG-AT1R, early endosomal targeting peptide rGFP-2xFYVE, and β-arrestin 1-RFP **(c)** or β-arrestin 2-RFP **(d)** pre-stimulation and after 45-minute stimulation of 1 μM AngII.

**Extended Data Figure 2:**
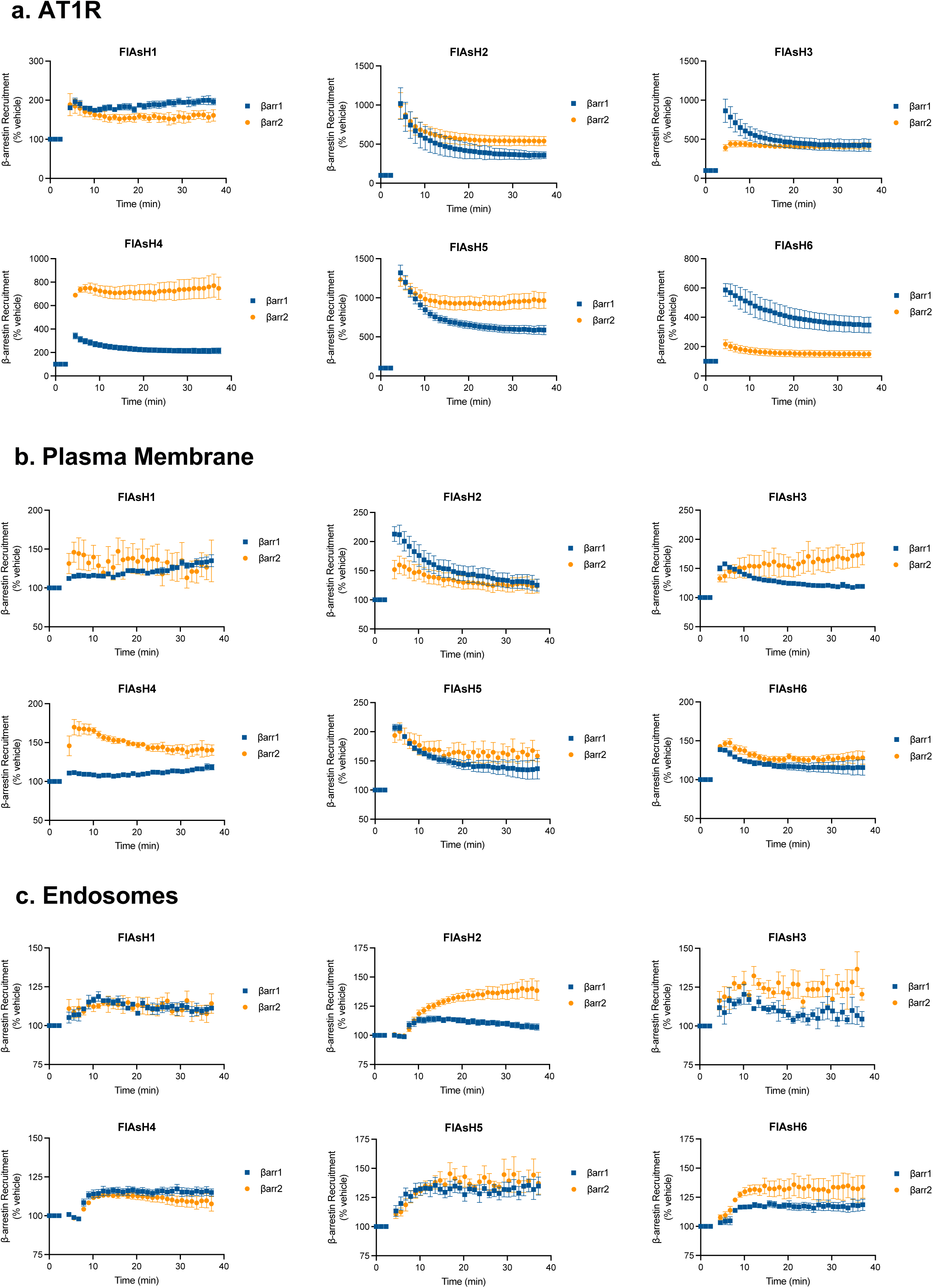
Recruitment of NanoBiT FlAsH biosensors to the AT1R, plasma membrane, and early endosomes. **(a)** For recruitment to AT1R, HEK293T cells were transfected with AT1R-LgBiT, one of the six SmBiT-β-arrestin1-FlAsH biosensors or six SmBiT-β-arrestin2-FlAsH biosensors, and stimulated with 1 μM AngII. **(b)** For recruitment to the PM, HEK293T cells were transfected with FLAG-AT1R, LgBiT-CAAX, and SmBiT-β-arrestin-FlAsH biosensors and stimulated with 1 μM AngII. **(c)** For recruitment to endosomes, HEK293T cells were transfected with FLAG-AT1R, 2xFYVE-LgBiT, and SmBiT-β-arrestin-FlAsH biosensors and stimulated with 1 μM AngII. The luminescence signal from the complementation of NanoBiT fragments was normalized to pre-stimulation signals and then normalized to vehicle. Data represents mean ± SEM of *n* independent biological replicates, n=4 for CAAX and AT1R, n=5 for 2xFYVE.

**Extended Data Figure 3:**
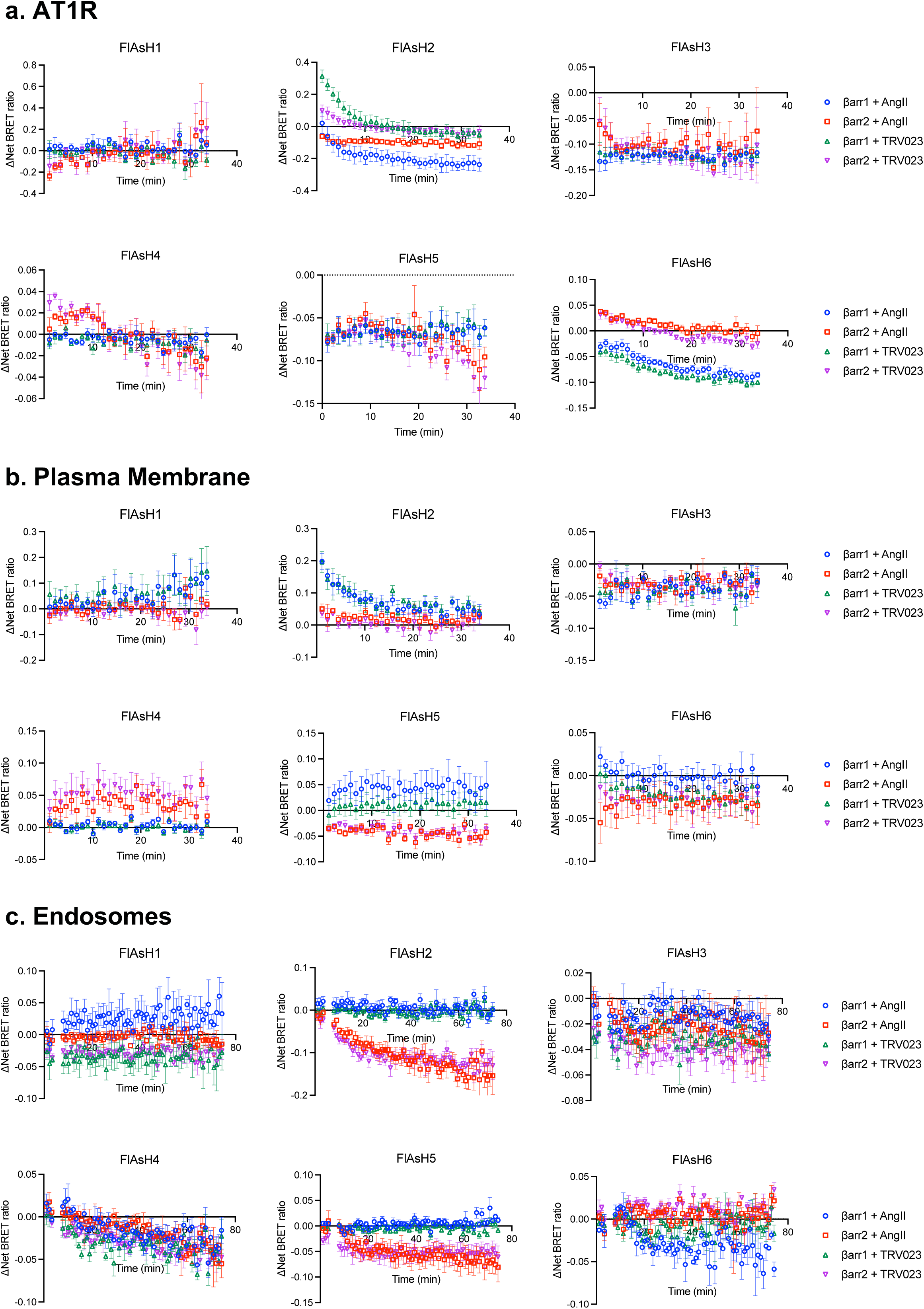
Kinetic tracings of BRET signals from NanoBiT FlAsH assays to detect location-specific β-arrestin conformations. **(a)** For β-arrestin conformations at AT1R, HEK293T cells were transfected with AT1R-LgBiT and one of the six SmBiT-β-arrestin1-FlAsH biosensors or six SmBiT-β-arrestin2-FlAsH biosensors. **(b)** For β-arrestin conformations at the PM, HEK293T cells were transfected with FLAG-AT1R, LgBiT-CAAX, and SmBiT-β-arrestin-FlAsH biosensors. **(c)** For β-arrestin conformations in endosomes, HEK293T cells were transfected with FLAG-AT1R, 2xFYVE-LgBiT, and SmBiT-β-arrestin-FlAsH biosensors. Cells were then labeled with the arsenic dye FlAsH-EDT2 or HBSS mock label and stimulated with 1 μM AngII or 10 μM TRV023. ΔNet BRET ratio was calculated by subtracting the net BRET values of FlAsH-labeled cells from the mock-labeled condition. Data represents mean ± SEM of *n* independent biological replicates. For AT1R, FlAsH 1: n=3, FlAsH 2,4: n=4, FlAsH 3, 5, 6: n=5. For CAAX, FlAsH 1-4: n=4, FlAsH 5: n=5, FlAsH 6: n=6. For 2xFYVE, FlAsH 1-5: n=4, FlAsH 6: n=3. Data with TRV023 has the same number of replicates as AngII, except FlAsH 5 (CAAX): n=7 and FlAsH 2 (FYVE): n=5.

**Extended Data Figure 4:**
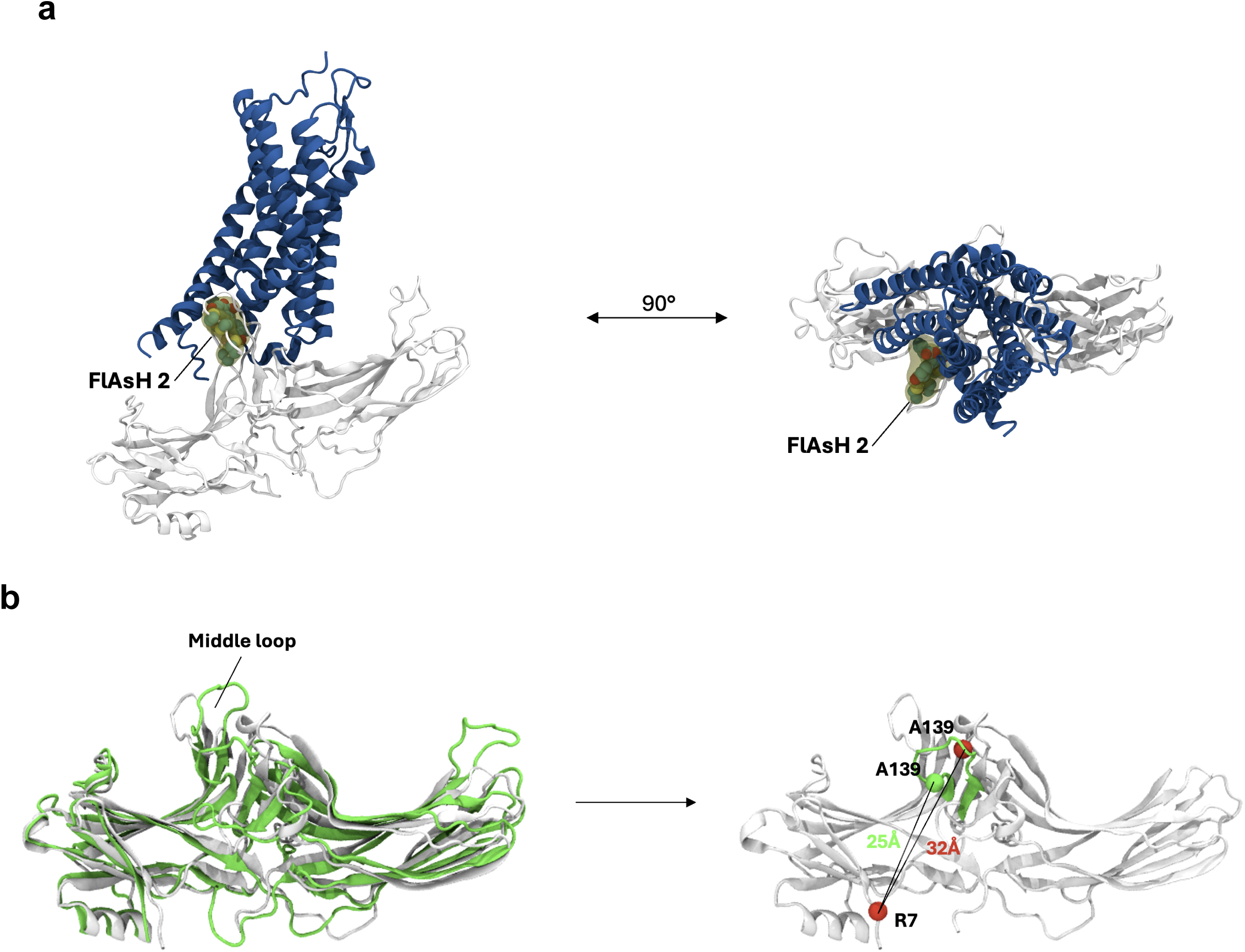
Orientation of FlAsH 2 in different configurations of receptor-β-arrestin1 complex. **(a)** Spatial orientation of FlAsH 2 in a β-arrestin 1-GPCR core complex. Using the structure of the neurotensin receptor 1 (blue) in complex with β-arrestin 1 (gray) (PDB: 6UP7), we manually modelled the full-length FlAsH 2 into the complex (yellow). Subsequently, the structure was subjected to a short minimization run (0.1 RMS kcal/mol/Å^2^ gradient, Amber10:EHT forcefield). The resulting structural model suggests that in the GPCR core complex, FlAsH 2 would intercalate into the receptor structure, thus reducing its mobility and resulting in a low BRET ratio. **(b)** Spatial orientation of FlAsH 2 in a β-arrestin 1-GPCR C-tail complex. Comparison of the structures of the V2Rpp-bound (green, PDB: 4JQI) and inactive (white, PDB: 1G4M) β-arrestin 1. The position of the insertion of FlAsH 2 (A139) within the middle loop is depicted as a sphere (green – 4JQI, red – 1G4M). The distance between FlAsH 2 and the NLuc donor was approximated by plotting the distance between the insertion position and the N-terminally located R7.

**Extended Data Figure 5:**
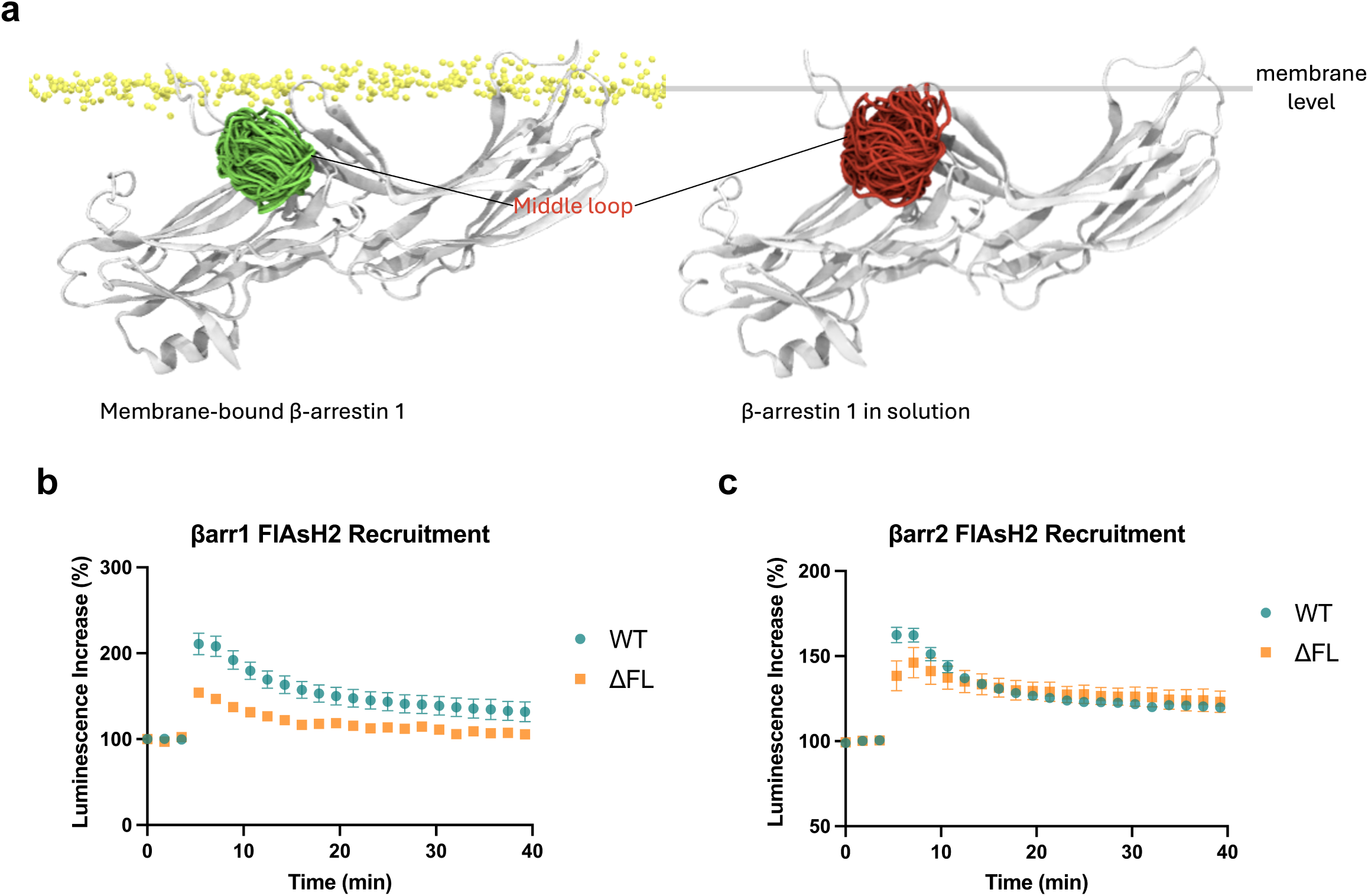
Membrane insertion of the finger loop affects the middle loop’s orientation. **(a)** MD simulations comparing the orientation of the middle loop of membrane-anchored β-arrestin 1 vs β-arrestin 1 in solution. The flexibility of the middle loop was illustrated using structural snapshots from accumulated frames (one snapshot every 30 ns) for β-arrestin 1 embedded in membrane (green) and β-arrestin 1 in solution (red). **(b)** Recruitment of FlAsH 2 biosensors for WT β-arrestins and finger loop deletion mutants (ΔFL) to the PM using the NanoBiT assay. Data shown represents mean ± SEM, n=4 independent biological replicates.

**Extended Data Figure 6:**
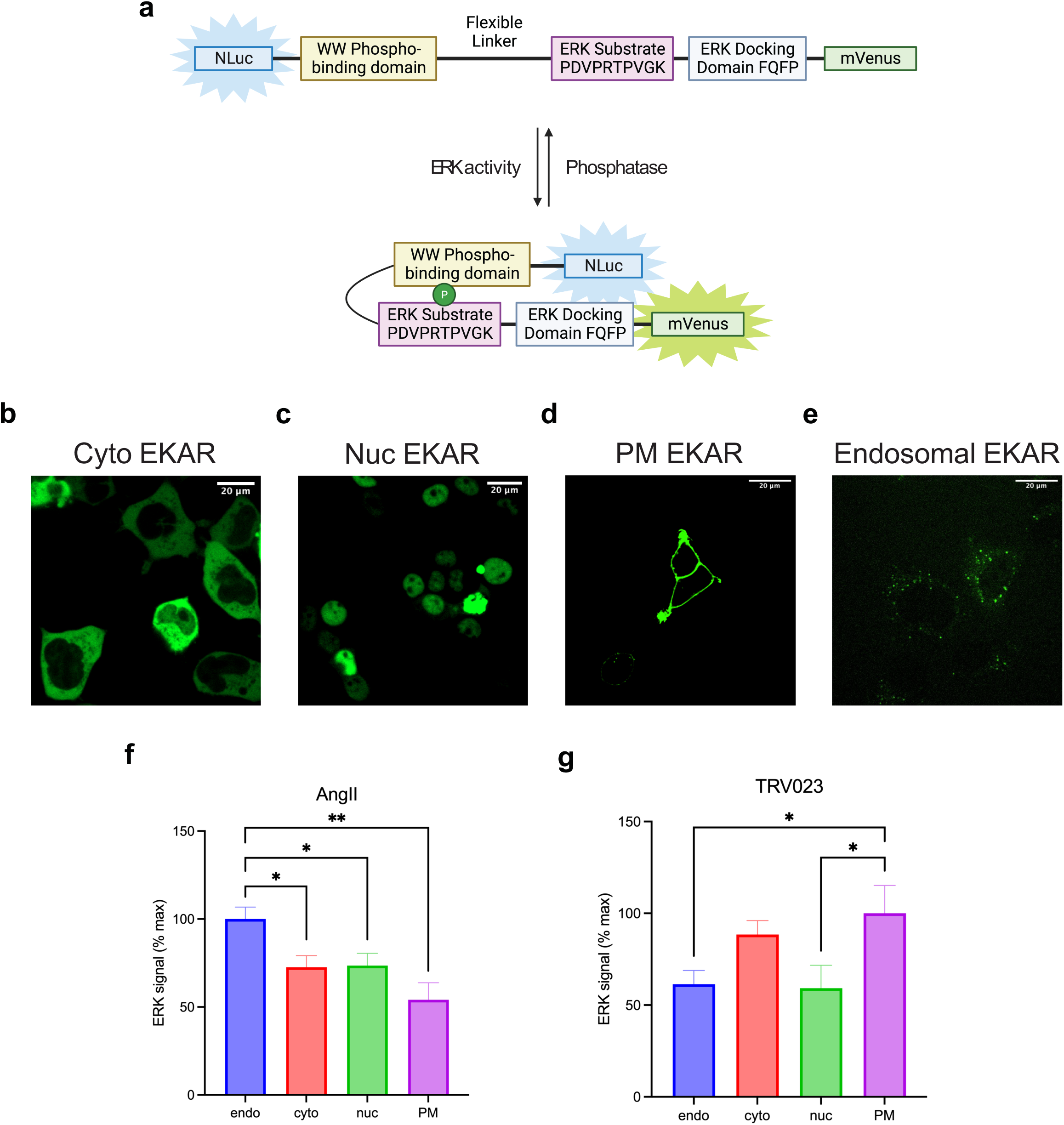
EKAR BRET biosensors to assess location-specific ERK1/2 activation in HEK293T cells. **(a)** Schematic of BRET-based EKAR biosensors adapted from the previously published FRET versions^58^ (**b-e**) Confocal microscopy images of EKAR biosensor expression in the cytosol, nucleus, PM, and early endosomes. **(f-g)** Distinct distribution of subcellular pools of ERK signaling promoted by AngII and TRV023. Data was quantified as AUC of BRET signals over 50 minutes after ligand stimulation and normalized to the max signal of each ligand. Data represents mean ± SEM of *n* independent biological replicates, n=4 for PM and cytosolic ERK, n=5 for nuclear and endosomal ERK. One-way ANOVA with Holm-Šídák’s posthoc test comparing to subcellular location with max signal. *P<0.05; **P<0.005.

**Extended Data Figure 7:**
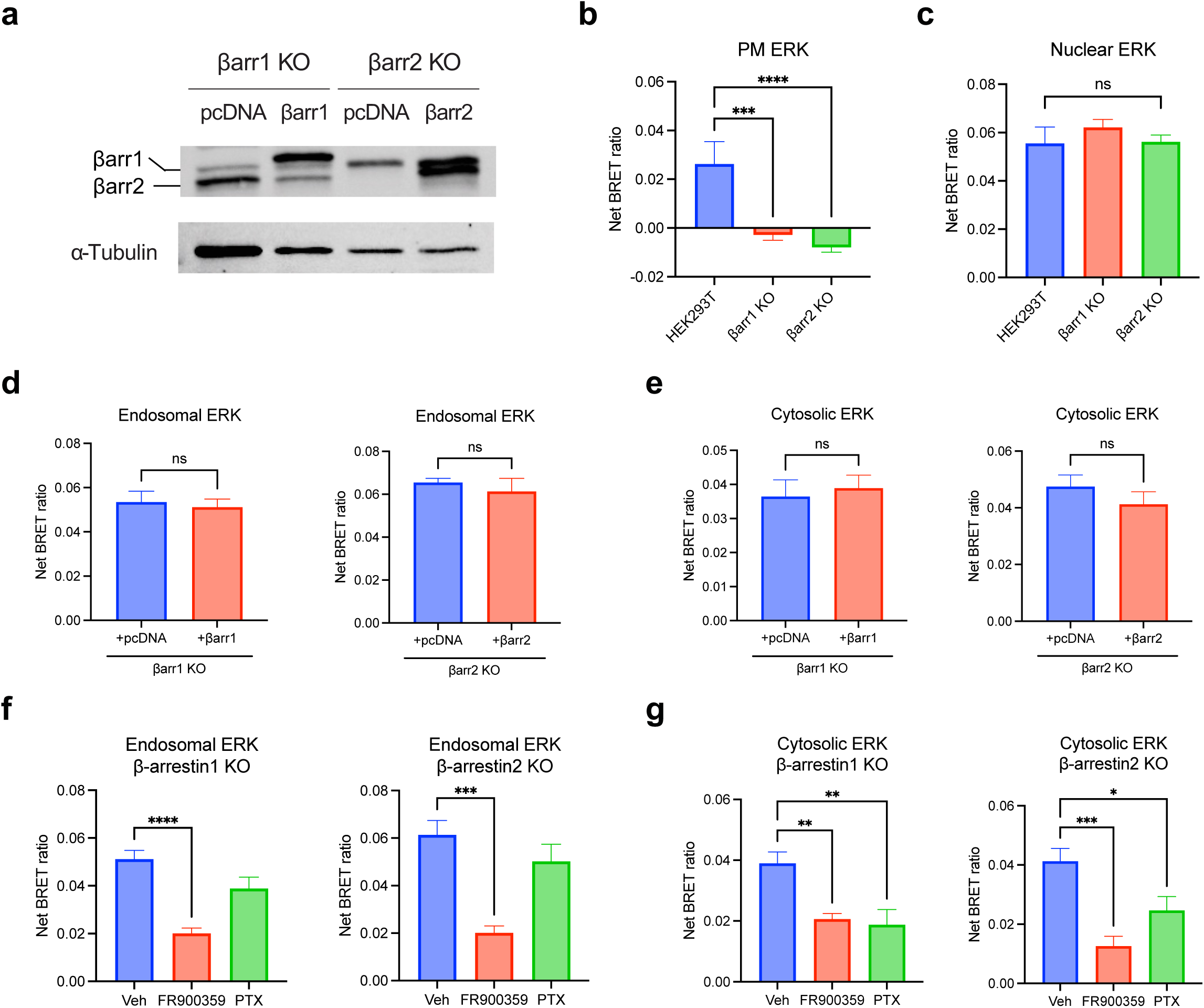
EKAR BRET biosensors to assess location-specific ERK1/2 activation in β-arrestin KO HEK293 cells. **(a)** Representative western blot of β-arrestin1/2 (A1CT) in β-arrestin 1 KO and β-arrestin 2 KO HEK293 cells with pcDNA control or FLAG-β-arrestin rescue. n=3 independent biological replicates. **(b, c)** PM and nuclear ERK activity in WT HE293T, β-arrestin 1 KO, and β-arrestin 2 KO cells. Data represents mean ± SEM of *n* independent biological replicates, n=5 for HEK293T, n=8 for β-arrestin 1 and 2 KO cells. One-way ANOVA with Dunnett’s multiple comparison test to compare β-arrestin KO cells vs HEK293T. ***P<0.0005; ****P<0.0001; ns, not significant. **(d, e)** Endosomal and cytosolic ERK activity in β-arrestin 1 or β-arrestin 2 KO HEK293 cells upon rescue with pcDNA control or FLAG-β-arrestin 1 or FLAG-β-arrestin 2. Cells were stimulated for 30 minutes with 1 μM AngII. Data represents mean ± SEM, n=7 independent biological replicates. Unpaired two-tailed t-tests comparing pcDNA vs. β-arrestin rescue. ns, not significant. **(f, g)** Effect of Gq inhibition and Gi inhibition using FR900359 and PTX, respectively, on endosomal and cytosolic ERK signaling in β-arrestin 1 or β-arrestin 2 KO HEK293 cells overexpressing FLAG-βarrestin 1 or FLAG-βarrestin 2. Data represents mean ± SEM, n=7 independent biological replicates. One-way ANOVA with Šídák’s posthoc test comparing inhibitors vs vehicle. *P<0.05; **P<0.005; ***P<0.0005; ****P<0.0001.

## Notes

### Competing Interest Statement

The authors have declared no competing interest.

